# A platform for deep topographic multiomic mapping of the bone marrow environment in multiple myeloma

**DOI:** 10.64898/2026.09.22.753531

**Authors:** Jinsen Lu, Chen-Yi Wang, Simon Davis, Edmund Watson, Chao Jiang, Warren Baker, Eleanor Calcutt, Sarah Gooding, Joanna Hester, Fadi Issa, Patricia Benevent Bellver, Alessandro Lagana, Chad Bjorklund, Giorgio Napolitani, Sue Griffin, Anjan Thakurta, Karthik Ramasamy, Anna Sozanska, Rosalin Cooper, Dan Royston, Nick Athanasou, Adam P Cribbs, Roman Fischer, Udo Oppermann, Srinivasa Rao

**Author notes:** Equal contributions (Jinsen Lu, Chen-Yi Wang, Simon Davis, Roman Fischer, Udo Oppermann, Srinivasa Rao).

## Abstract

Multiple myeloma is a genetically complex plasma cell malignancy in which profound intra-tumoral heterogeneity shapes disease progression and therapeutic resistance. Genomic subclones dynamically emerge and evolve under treatment pressure, yet their spatial organization within intact bone marrow remains poorly defined. We hypothesized that genetically distinct myeloma subclones occupy spatially discrete ecological niches associated with unique molecular and microenvironmental states. To test this, we here demonstrate a proof-of-concept spatial tri-omics workflow combining laser capture microdissection-based genomics and deep LC-MS/MS proteomics with single-cell spatial transcriptomics, integrated by digital co-registration of adjacent tissue sections. Applied to a myeloma bone marrow trephine, this approach enabled coordinated profiling of matched anatomical regions demonstrating that genomic, transcriptomic and proteomic profiles converged to define clone-specific microenvironmental programmes.

We show that genetically distinct myeloma subclones occupy mutually exclusive regions of the same bone marrow biopsy. Remarkably, an aggressive del(17p) clone and a gain(12q) clone occupied defined anatomical territories within the same trephine-biopsied section, each clone exhibiting distinct genomic, transcriptomic, proteomic and cellular ecosystems. The del(17p) clone localized to a fibrotic, osteolytic and angiogenic niche enriched for activated stromal cells, osteoblasts, osteoclasts and immunoregulatory myeloid populations, whereas the gain(12q) clone occupied a comparatively quiescent marrow niche with limited stromal remodelling. Together, these findings demonstrate that genetically distinct myeloma subclones establish spatially and biologically discrete ecological niches through coordinated remodelling of stromal, immune and bone compartments, providing a framework for understanding clonal evolution within the bone marrow microenvironment.

## INTRODUCTION

The emergence of spatially resolved molecular technologies has transformed the study of complex tissues by enabling the interrogation of gene expression, protein abundance, and genomic alterations within their native architectural context ^1–5^. Bulk sequencing approaches obscure cellular heterogeneity by averaging signals across mixed populations, while single-cell technologies, although providing high-resolution molecular insights, inherently disrupt spatial relationships that are critical for understanding tissue organization and cell–cell interactions. Spatial transcriptomics bridges this gap by retaining positional information of gene expression within intact tissue sections. This enables the mapping of cellular niches, tumour–immune interactions, and microenvironmental gradients that drive disease progression and therapeutic resistance. In oncology and inflammatory diseases, where tissue architecture and cellular crosstalk are fundamental determinants of pathogenesis, spatial approaches have rapidly become indispensable tools for dissecting disease biology.

Recent work has extended beyond single-modality spatial profiling toward multimodal spatial omics, integrating transcriptomics with proteomics, metabolomics, epigenomics, and imaging data ^6, 7^. Such approaches enable simultaneous characterization of multiple molecular layers within the same tissue section, offering a more comprehensive view of cellular states and interactions. Combined spatial transcriptomics and mass spectrometry imaging, for example, has demonstrated the feasibility of co-mapping RNA and metabolite distributions, overcoming previous technical incompatibilities between platforms^8^. More broadly, spatial multi-omics strategies have been shown to reveal complex intra-and intercellular molecular networks and to enhance the identification of disease-relevant pathways and therapeutic targets. To the best of our knowledge, however, no study has combined whole-genome sequencing, unbiased liquid chromatography-tandem mass spectrometry (LC-MS/MS) proteomics and spatial transcriptomics from spatially matched regions of the same tissue.

In parallel, multimodal integration is also advancing in the single-cell domain, where transcriptomic, proteomic, and chromatin accessibility data are increasingly combined to define cell states with greater precision ^9, 10^. However, these approaches still lack spatial context, necessitating computational or experimental strategies to align single-cell data with spatial frameworks. Consequently, there is a growing trend toward integrated spatial–single-cell pipelines, in which single-cell sequencing and immune profiling are used to deconvolute spatial signals, validate cell-type assignments, and infer functional interactions within tissue niches^11^. This convergence of technologies represents the current state of the art in systems-level tissue analysis and underpins the development of next-generation precision medicine approaches.

Multiple myeloma (MM) represents a compelling disease context in which to apply such integrated spatial and single-cell methodologies. MM is a hematologic malignancy characterized by the clonal proliferation of plasma cells within the bone marrow, leading to the overproduction of monoclonal immunoglobulins and subsequent end-organ damage^12, 13^. The disease evolves through a multistep process, typically progressing from monoclonal gammopathy of undetermined significance (MGUS) to smouldering myeloma and ultimately to symptomatic MM ^12–14^.

A defining feature of MM is its profound spatial and cellular heterogeneity within the bone marrow microenvironment. Malignant plasma cells coexist with diverse stromal, immune, and endothelial cell populations, forming a complex ecosystem that supports tumour growth, immune evasion, and drug resistance ^13, 15^. Interactions between myeloma cells and the bone marrow niche - mediated through cytokines, cell-cell contact, and extracellular matrix components - drive key pathological processes, including osteolytic bone disease, angiogenesis, and immunosuppression ^16, 17^.

Importantly, this microenvironmental complexity is not adequately captured by bulk or even single-cell sequencing alone. Spatial context is critical for understanding the localization of malignant clones, the distribution of immune infiltrates, and the formation of protective niches that contribute to relapse and treatment failure. Given that MM remains incurable in most patients despite significant therapeutic advances, there is a pressing need for technologies that can resolve these spatially organized interactions at multiple molecular levels.

In this proof-of-concept study, we address this need by applying an integrated spatial multi-omics approach to a relapsed patient MM bone marrow biopsy. By combining spatial transcriptomics, laser capture microdissection based unbiased proteomics via LC-MS/MS, and spatial genomics through a customized DNA sequencing strategy, we demonstrate a comprehensive, spatially resolved molecular map of the tumour microenvironment. Furthermore, we use single-cell sequencing and immune profiling to corroborate and refine spatial findings, enabling robust cross-modality validation.

## RESULTS

### Development of a spatial multi-omics platform

We established a spatial multi-omics workflow by integrating laser capture microdissection (LCM) based genomics and proteomics with in-situ targeted transcriptomics. We applied this platform approach to a human multiple myeloma FFPE bone marrow biopsy sample **(Fig. 1).** The sample was obtained from a 59-year-old male patient, diagnosed with kappa light chain myeloma and undergoing multiple lines of prior therapies including stem cell transplant, combinations of dexamethasone, proteasome inhibitors and immunomodulatory drugs including mezigdomide as part of a clinical trial, but no BCMA-targeted therapies. **(SI Fig 1)**. Cytogenetic analysis by fluorescence in situ hybridisation (FISH) as part of routine clinical bone marrow investigations suggested high risk features including gain (1q21) (22% of plasma cells), chr 14 rearrangements and del 17p13 (13% of plasma cells).

**Figure 1:**
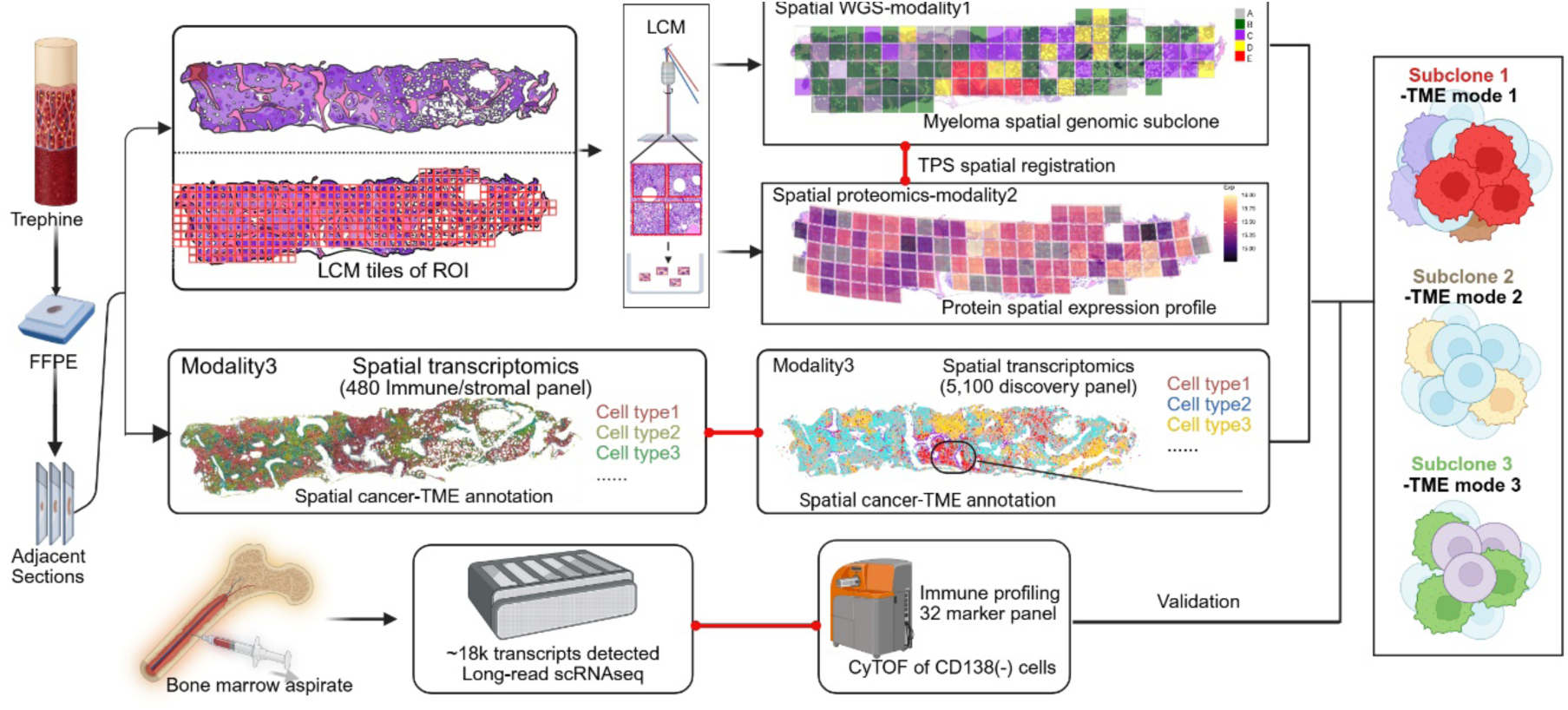
Overview of a spatial multiomics workflow to study myeloma clonality and tumour microenvironments (TME). Adjacent FFPE sections from a myeloma bone marrow trephine were tiled into 250 micrometre squares and subjected to laser capture microscopy (LCM) followed by whole genome sequencing and LC-MS/MS based deep proteomics. Other sections were subjected to spatial transcriptomics (Xenium, 480 and 5100 marker panels). Bone marrow aspirates from the same patient were subjected to long-read single cell RNAseq and mass cytometry to corroborate spatial results.

Consecutive 3-5 µm tissue sections were cut from the trephine, then assigned to different modalities including *in situ* spatial transcriptomics using customized cancer-immune marker panels on the Xenium platform with two custom marker panels (480 and 5.1k) **(SI File 1)**. Two subsequent sections were segmented into 250 µm tiles using LCM (404 tiles for the WGS layer; 384 tiles for the proteomics layer). Sets of four adjacent tiles were pooled during collection, yielding 101 merged tiles for WGS and 96 for proteomics, which were processed using the Adaptive Resolution Multiscale Spatial (ARMS) DNAseq workflow ^18^ for shallow whole-genome sequencing (sWGS) and for liquid chromatography-trapped ion mobility spectrometry (LC-TIMS/TOF) based 4D proteomics^19^. A digital registration workflow using the thin plate spline (TPS) method^20^ computationally aligned the Xenium data with the LCM tile coordinates from the adjacent sections, correcting for non-linear distortion and enabling multi-modal analysis of matched tissue areas of serial sections. This integrated platform yielded registered, three-layer data across the entire bone trephine (∼1.5 cm×0.3 cm).

### Identification of myeloma clonal structures by spatial WGS

A copy number alteration (CNA) profile was derived from the tiled LCM-WGS data **(Fig. 2A, SI File 2),** leading to identification of distinct tumour containing regions (‘A’-‘E’) across the trephine (Copykit 0.1.4). Phylogenetic analysis of the tile sections identified region ‘A’ with low tumour purity, whereas region ‘B’ suggests an ancestral subclone ‘B’ which harboured a set of basal CNA features (gain: chr6, chr7q, chr9q, chr17q, chr18 and chr19; loss: chr12p, chr13 and chr14). The ‘B’ lineage then branched into two, with subclone ‘C’, characterized by an additional gain of chr12q. Subclone ‘B’ also developed into ‘D/E’ including loss of chr2q, gain of chr9p and chr15. Specifically, subclone ‘E’ acquired loss of chr17p and gain of 5q. These subclonal genomic CNA profiles were further corroborated by pseudobulk CNA profiles from ARMS DNAseq as well as profiles inferred from spatial bulk-RNA as well as from long-read scRNA-seq data (Copykat v1.2.5), obtained from the same patient’s contemporaneous bone marrow aspirate **(Figs. 2B and 2C, SI Fig. 2, SI Fig. 3)**.

**Figure 2:**
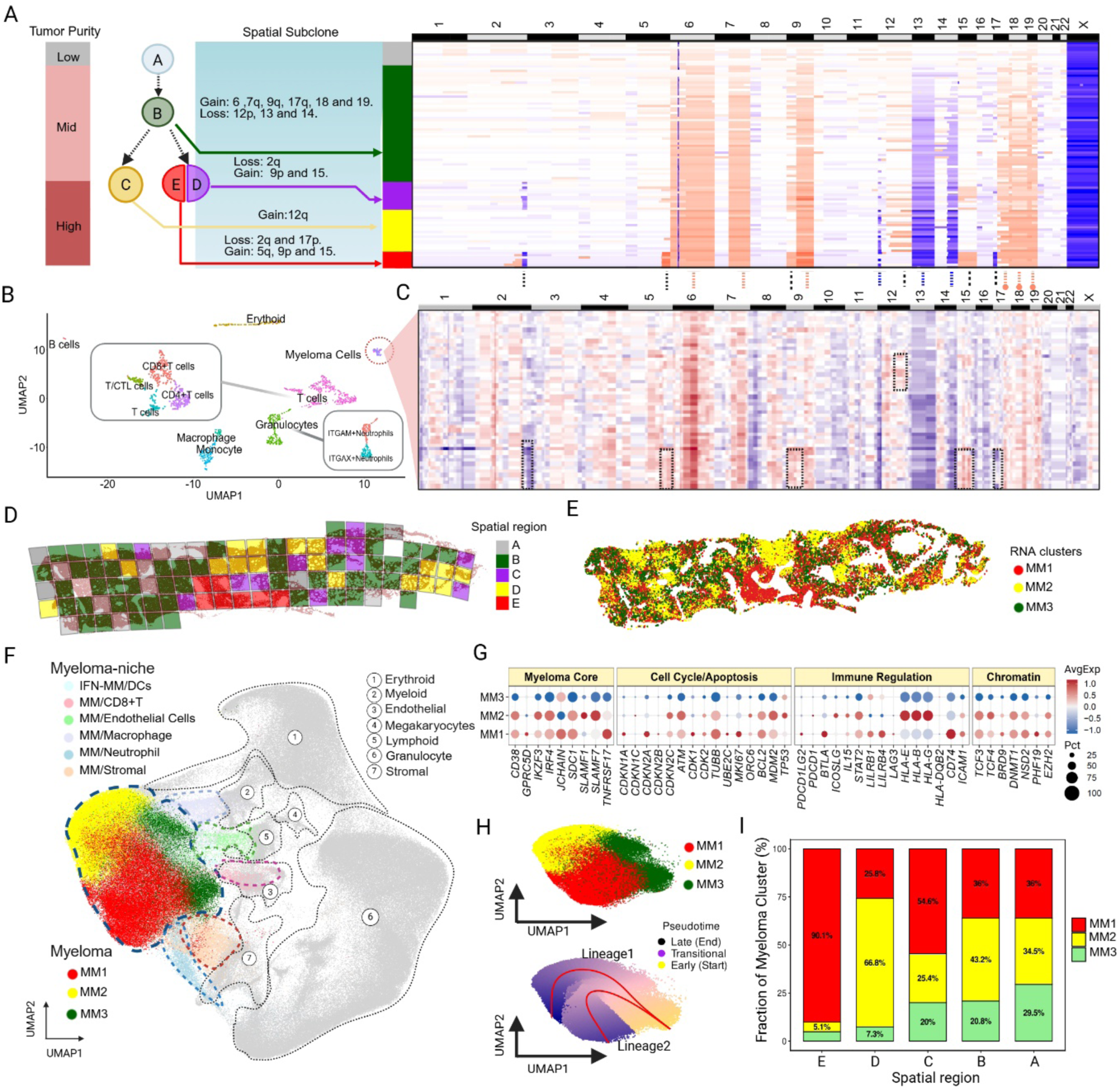
Subclonal structure and spatial distribution identified by shallow WGS. **(A)** Identification of subclonal regions by laser-capture microscopy (LCM) mediated whole genome sequencing (WGS) of individual LCM tiles resulting in spatial genomic regions ‘A’-‘E’ through copy number alteration (CNA) profiles from WGS data. (**B)** UMAP of major cell clusters from scRNA data from bone marrow aspirates of the same patient. (**C** ) Heatmap of CNA profile computationally inferred from the myeloma population identified in scRNA data from a bone marrow aspirate of the same patient. (**D)** Spatial map of subclone regions ‘A’-‘E’. (**E)** Distribution of myeloma RNA clusters MM1-MM3 across a subsequent section, identified through their spatial transcriptomic profile. (**F)** UMAP of spatial transcriptomic clusters and cell types (480 gene panel). Major myeloma clusters (MM1-red, MM2-yellow, MM3-green) highlighted. (**G)** Dotplot of canonical markers of myeloma, cell cycle, immune and chromatin factors for MM1-MM3. (**H)** Pseudotime analysis of myeloma clusters MM1-MM3. (**I)** Distribution of myeloma clusters MM1-MM3 in WGS derived spatial subclone regions.

### Myeloma subclones align with transcriptionally distinct myeloma clusters

We investigated the trephine cell type composition by deploying a custom 480 gene panel designed towards identification of myeloma-immune-stromal interactions using the Xenium spatial transcriptomics platform. The trephine section used for this spatial transcriptomics experiment comprises ∼301,000 cells, with a malignant plasma cell content of ∼30%. We identified 8 major cell types, including 3 distinct myeloma clusters (MM1-MM3), while deeper clustering revealed 33 distinct cell types and environmental niches (**SI Fig. 4**). Comparing the spatial distribution between genomic tiles and transcriptional myeloma clusters reveals an excellent overlap (MM1 corresponding to ‘D/E’, del 17p; MM2 corresponding to ‘C’ gain 12q; MM3 corresponding to truncal subclone ‘B’) **(Figs 2D-2E)**. Gene expression in the two main myeloma clusters mirrored these copy number changes. Compared with MM2, MM1 expressed lower TP53, located on the deleted chr17p, and higher PDCD1LG2 (PD-L2), which lies on the gained chr9p. MM2 expressed higher MDM2, which lies on the gained chr12q (**Fig. 2G, SI File 7**). Pseudotime analysis of MM1-MM3 gene transcription profiles **(Fig 2H)** reveals a trajectory emanating from cluster MM3 and branching into 2 distinct branches (MM1 and MM2, respectively), in accordance with the identified genomic subclonal hierarchy **(Fig 2A).** Furthermore, aligning the genomic tile regions with the Xenium-derived spatial pattern of myeloma cell distributions identified areas of distinct myeloma cluster occupations **(Fig 2I)**. In particular, spatial region (as determined by LCM/WGS) ‘E’ is predominantly dominated by myeloma cluster MM1 (90.1 %), while region ‘D’ is largely occupied by MM2 (66.8 %).

### Multiple immunosuppressive mechanisms restrict cytotoxic responses within an inflamed myeloma microenvironment

To gain further insights into immune phenotypes we investigated an adjacent section using a custom 5100 marker Xenium panel **(SI File 1)**. This resulted in identification of 36 cell types **(Fig, 3 and SI Fig 5)** with good overlap between cell types identified by the 480 panel Xenium experiment (**SI Fig. 4**). Spatial transcriptomic analysis identified an immune-infiltrated myeloma microenvironment **(Fig. 3A-B)** containing three distinct cytotoxic lymphocyte populations and two tumour-associated macrophage (TAM) populations. All myeloma clusters expressed the stress-induced NKG2D ligands MICB and, to a lesser extent, MICA, **(Fig. 3C-D)** suggesting that malignant plasma cells retain features capable of promoting NK-cell and cytotoxic T-cell recognition.

**Figure 3:**
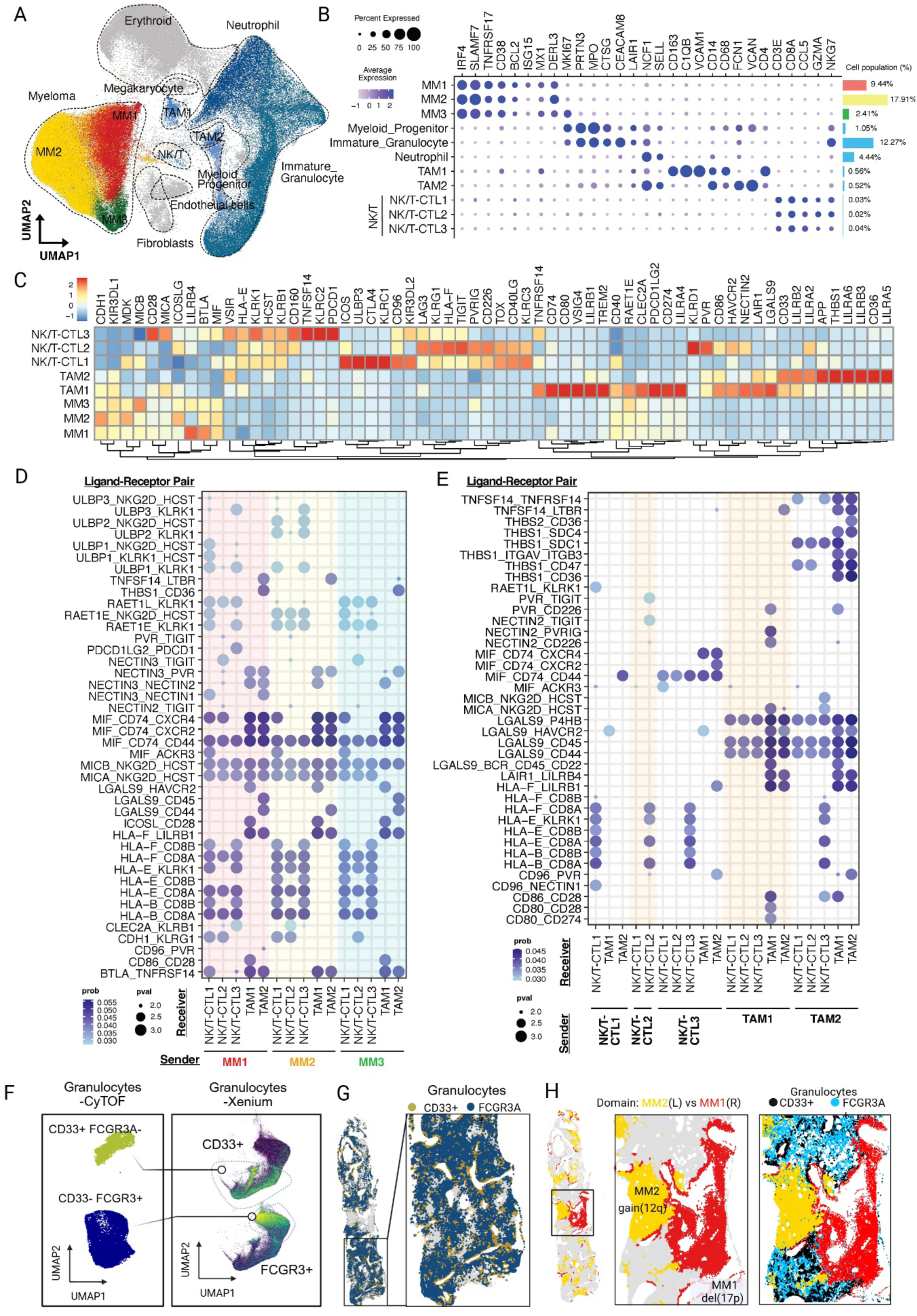
Spatial proteomics reveals an inflamed and immuno-suppressed environment. **(A)** UMAP from spatial transcriptomics (Xenium, 5.1k panel) showing myeloma clusters (MM1-MM3) and major bone marrow populations. **(B)** Dotplot illustrating marker genes and relative proportions of selected immune cell clusters. (**C)** Heatmap highlighting expression levels of selected markers for immune interactions between myeloma, lymphoid and tumour associated macrophage clusters. (**D)** Dotplot of CellChat analysis for immune interactions with myeloma clusters as Sender. (**E)** Dotplot of CellChat analysis for interactions between cytotoxic lymphoid and immunosuppressive macrophage clusters. (**F)** UMAPs derived from mass cytometry (bone marrow aspirate of the same patient) and spatial transcriptomic showing marker differences (CD33, FCGR3A) for two granulocyte clusters. (**G)** CD33+ and FCGR3A+ granulocytes show distinct localisation across the trephine section with CD33+ cells showing selective endosteal localisation. (**H)** Distribution of myeloma clusters MM1 and MM2 across the trephine. Both CD33+ and FCGR3A+ granulocytes (dark and light blue, respectively) are localised at the periphery to focal tumour regions.

Correspondingly, all lymphocyte clusters expressed KLRK1 (NKG2D), indicating preserved capacity for tumour recognition through the NKG2D axis **(Fig. 3E)**. However, the cytotoxic compartment exhibited marked heterogeneity. NK/T-CTL1 cells displayed an activated, yet suppressive phenotype characterised by high KLRC1 (NKG2A), CD96, CTLA4, and TOX, consistent with chronic activation and emerging dysfunction. NK/T-CTL2 demonstrated the strongest exhausted phenotype, with high expression of TIGIT, LAG3, HAVCR2 (TIM-3), TOX, PVRIG, and KLRG1, while NK/T-CTL3 retained features of an activated effector population, including high KLRC2 (NKG2C), PDCD1, CD28, and KLRK1 **(Fig. 3C).** The tumor-associated macrophage (TAM) compartment **(Fig. 3C)** exhibited a predominantly immunoregulatory phenotype. TAM1 expressed high levels of PD-L1 (CD274), PD-L2 (PDCD1LG2), LILRB1, LILRB2, LGALS9, HAVCR2, TREM2 and LAIR1, consistent with a highly suppressive macrophage state. TAM2 shared expression of LILRB1, LILRB2, LAIR1, and CD86, although with lower expression of PD-L2, suggesting a less activated but still regulatory phenotype. Together with the concurrent expression of TIGIT, PDCD1, TIM-3, and PVRIG within cytotoxic lymphocytes, these findings indicate the presence of multiple convergent inhibitory pathways capable of restraining anti-tumour immunity.

### Non-classical HLA signalling contributes to immune suppression within an inflamed myeloma microenvironment

The expression of HLA-E by myeloma cells, TAMs, and lymphocyte populations **(Figs. 3C-E)** suggests an important role for non-classical MHC-mediated immune regulation. Given the prominent expression of NKG2A (KLRC1) within cytotoxic NK/T lymphocytes (**Fig. 3C),** HLA-E-mediated inhibitory signalling may suppress NK-cell cytotoxicity despite persistent MICA/MICB expression, representing a potential mechanism of immune escape in this niche **(Fig. 3D).** This interpretation is supported by studies demonstrating that HLA-E expression on myeloma cells impairs NKG2A-positive NK-cell function and is associated with immune dysfunction in myeloma ^21, 22^. In contrast, the high expression of NKG2C (KLRC2) within NK/T-CTL3 (**Fig. 3C**) suggests that responses to HLA-E may vary across cytotoxic subsets, potentially reflecting a balance between inhibitory and activating signals. Notably, HLA-F was highly expressed in activated lymphocyte populations and was accompanied by abundant expression of its reported receptors LILRB1, LILRB2, and KIR3DL2 within macrophage and lymphocyte compartments **(Fig. 3C-F)**. To our knowledge, the functional significance of HLA-F in multiple myeloma has not been investigated, however, HLA-F has been shown to interact with both KIR and LILRB family receptors and is increasingly recognised as a mediator of immune regulation in cancer ^23^. The coordinated expression of HLA-F and its cognate receptors in our dataset raises the possibility that non-classical HLA-dependent signalling contributes to macrophage-and lymphocyte-mediated immune suppression. Collectively, these data support a model in which the myeloma microenvironment remains immunologically active, as evidenced by preserved NKG2D ligand expression and infiltration by cytotoxic lymphocytes but is functionally constrained by overlapping checkpoint pathways. The simultaneous presence of HLA-E/NKG2A, PD-1/PD-L1/PD-L2, TIGIT/PVR, TIM-3/Galectin-9, and potentially HLA-F/LILRB/KIR signalling is consistent with an immune-inflamed but highly regulated tumour microenvironment in which anti-myeloma immune responses are actively restrained rather than absent.

### Granulocyte populations exhibit progressive acquisition of immunosuppressive features

**-**Spatial transcriptomic and immune mass cytometry analysis identified a continuum of granulocytic populations spanning granulocyte progenitors to mature neutrophils, consistent with active granulopoiesis within the bone marrow microenvironment **(SI Figs. 6A-C).** Rather than exhibiting a classical inflammatory phenotype, granulocytes acquired progressively immunoregulatory features during maturation, with sustained LAIR1 expression and increasing LILRB2, HLA-E, and PD-L1 expression in mature neutrophils, while MICA/B expression declined (**SI Fig. 6D**). This pattern suggests a shift from immature granulocytes potentially susceptible to NK-cell recognition toward mature neutrophils with enhanced capacity to suppress cytotoxic lymphocyte function through the HLA-E/NKG2A and PD-L1/PD-1 axes. In contrast to tumour-associated macrophages, granulocytes showed limited expression of canonical T-cell checkpoint molecules, indicating that their immunomodulatory activity is mediated primarily through inhibitory myeloid receptor pathways **(SI Fig. 6D)**. Mass cytometry of a CD138^-^fraction of the patient’s bone marrow aspirate corroborates the developmental trajectory highlighting a CD33+ early and FCGR3A/CD16a^+^ late granulocyte population **(Fig. 3F and SI Fig. 6F)**, with CD33+ granulocytes mainly located to the endosteal compartment **(Fig. 3G)**.

Granulocytes/neutrophils also demonstrated a pattern of exclusion from tumour-infiltrated areas, particularly in association with the MM1 and MM2 myeloma subclones **(Fig. 3H and SI Fig. 6E)**. Collectively, these findings support a role for differentiating granulocytes in establishing an immunosuppressive, immune-constrained myeloma microenvironment.

### Multi-omic integration reveals phenotypic and metabolic differences between trephine areas associated with distinct myeloma clones

A trephine section adjacent to that used for WGS-guided spatial genomics was LCM-tiled using a TPS-transformed grid and analysed by LC-TIMS/TOF. Out of the 96 merged tiles, 81 passed quality control, with 4,000–7,000 proteins identified per tile, providing deep spatial proteome coverage across the biopsy **(SI File 4)**. Following normalisation, unsupervised clustering resolved six spatial proteomic clusters. Comparison with spatial transcriptomic data, aggregated into matched tile-specific pseudo-bulk profiles, demonstrated strong concordance between transcriptomic and proteomic measurements across corresponding tile regions **(Figs. 4A-B)**. To test this concordance formally, we integrated the genomic, transcriptomic and proteomic layers of the 81 tiles that passed quality control in all three assays using Multi-Omics Factor Analysis (MOFA). Clustering of tiles on the MOFA factors reproduced the separately derived protein and RNA clusters (**SI Fig. 8C**).

**Figure 4:**
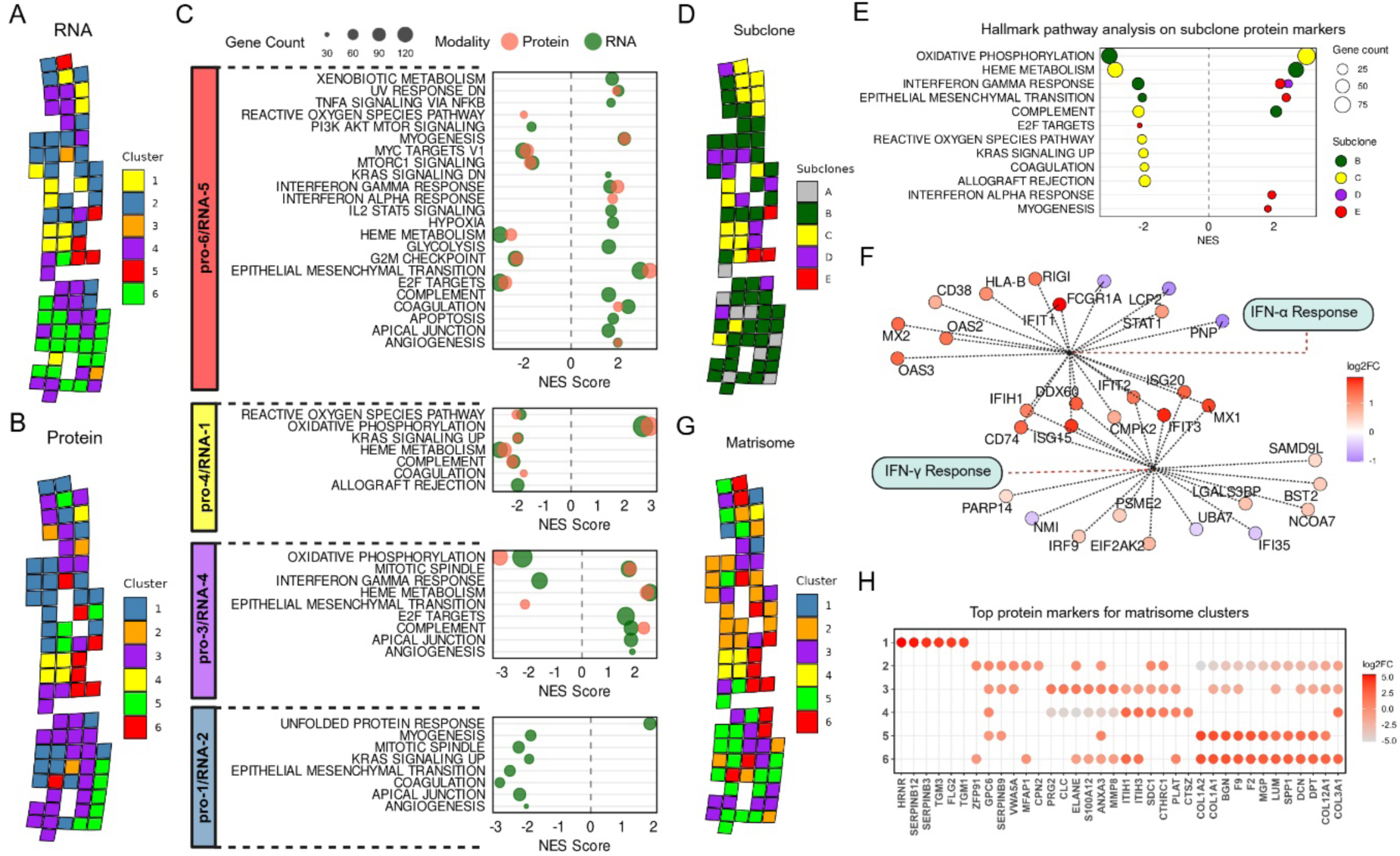
LC-MS/MS spatial proteomics identifies differences in metabolism and ECM composition between different subclone niches. **(A)** Tile-based clustering of pseudobulk RNA expression data from spatial transcriptomics. **(B)** Tile-based clustering of proteins derived from LCM based LC-MS/MS. **(C)** Pathway analysis overlapping of clusters (protein-RNA). **(D)** Tile-based clustering of genomic subclones from LCM based spatial genomic analysis. (E) Pathway analysis of subclonal regions. **(F)** IFN-α and IFN-γ response gene sets demonstrating enrichment in region of subclone E corresponding to MM1. (**G)** Tile-based clustering of matrisome proteins. (**H)** Dotplot of top differential matrisome proteins associated with matrisome clusters.

Matching clusters with their corresponding RNA and proteome profiles reveals significant differences between trephine regions. For example a matched region [pro-6/RNA-5] shows upregulation of angiogenesis, EMT and interferon responses, and downregulation of MTORC1 signalling, while a second matched region [pro-4/RNA-1] shows upregulation of oxidative phosphorylation **(Fig 4C).** Integration of subclone clusters **(Fig 4D, E)** with proteomic/transcriptomic data shows enrichment of interferon-α and interferon-γ responses in the MM1 related cluster region (protein cluster 6, RNA cluster 5 and subclone D/E, del(17p)). This enrichment was largely driven by the robust upregulation of classical interferon-stimulated genes, notably IFIT1, IFIT3, DDX60, and MX1 (**Fig 4F**), consistent with a recent single cell study associating type I IFN response with del(17p) myeloma ^24^. The same interferon programme emerged independently from the MOFA analysis. The proteins that most strongly marked the subclone D/E direction of Factor 4 were dominated by interferon-stimulated proteins, including ISG15, ISG20, MX1, MX2, OAS2, OAS3, IFIT3 and RIGI (**SI Fig. 8G**).

In addition, the identified geneset term “epithelial mesenchymal transition”, characterized by the pronounced expression of key matrix remodelling and adhesion molecules including POSTN, COL3A1, VCAN, and PCOLCE, and angiogenesis pathway proteins was enriched in MM1 relative to the MM2 related cluster region (protein cluster 4, RNA cluster 1 and subclone C) **(Fig 4C and E**), whereas MM2 demonstrated prominent enrichment of oxidative phosphorylation. For the erythroid and granulocyte dominant region (protein cluster 1/3, RNA cluster 2/4 and subclone B), pathway enrichment of heme metabolism, mitotic spindle, and unfolded protein response indicates cell division as well as a metabolically active, and cellularly stressed environment (**Fig 4C and E**). A spatial ECM tile profile (**Fig 4 G**) was derived from Matrisome-annotated proteins^25^ and tightly mirrored the global proteome clustering **(Fig 4B)**. The MM1 region (Matrisome cluster 6) exhibited a stroma-rich, fibrotic microenvironment enriched in fibrillar collagens (COL1A1, COL3A1) and proteoglycans (BGN, DCN). In contrast, the highly cellular MM2 region (Matrisome cluster 4) retained SDC1 but lacked structural collagens. Finally, adjacent erythroid and granulocyte-rich domains (Matrisome clusters 2 and 3) were defined by myeloid-derived proteases, including ELANE and MMP8 (**Fig 4H**). These findings demonstrate that spatial proteomics faithfully recapitulates genomic and transcriptomic tumour architecture while providing complementary information on metabolic state and extracellular matrix organisation within genetically distinct myeloma clones.

### Spatially resolved multi-omic profiling reveals clone-specific environmental niches in multiple myeloma

Integrated spatial transcriptomic, proteomic and genomic profiling demonstrated marked spatial segregation of MM1 (del17p) and MM2 (gain 12q) clones in specific regions located in the central part of the trephine **(Fig. 5A)** with minimal overlap between the two populations. This genomic compartmentalisation was accompanied by striking differences in tissue architecture and cellular composition, indicating that each clone occupies a distinct microenvironmental niche. Moreover, myeloma cells from MM1 and MM2 regions are distinguishable by their morphology in HE stains **(Fig. 5B)**.

**Figure 5:**
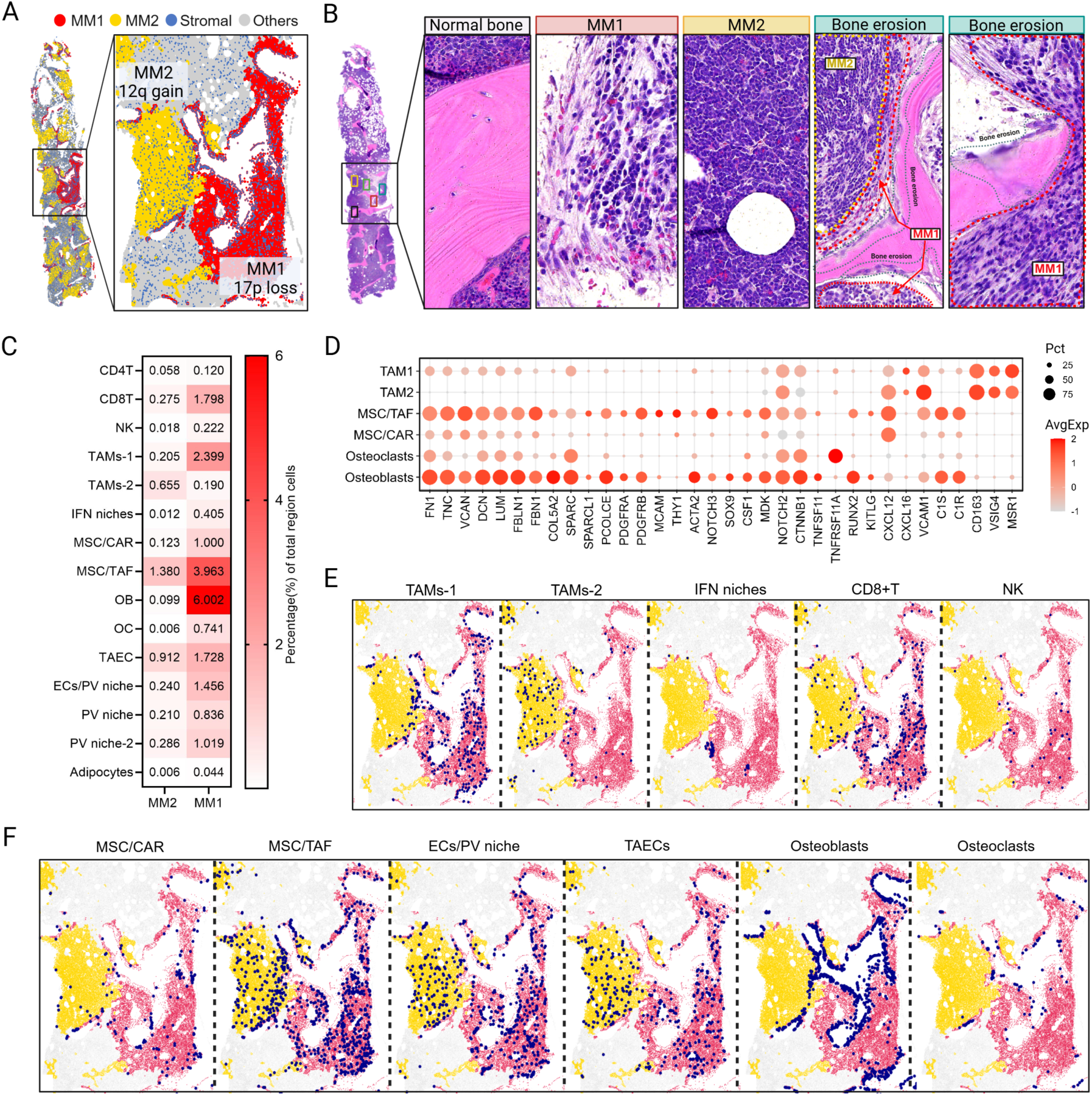
Myeloma subclones MM1 and MM2 are associated with fundamentally different TME niches and osteolytic and profibrotic environments. **(A)** Distribution of MM1 and MM2 subclones across the trephine section with central domains selected for subsequent analysis highlighted. (**B)** HE staining of trephine section areas. Magnifications show osteolytic trabecular bone lesions are exclusively associated to MM1 (del (17p)) but not to MM2 (gain (12q)). Morphology differences between MM1 and MM2 are depicted. (**C**) Heatmap showing the proportion of selected stromal and immune populations in MM1 and MM2 regions. (**D)** Dotplot of pro-fibrotic and osteolytic genes between TAM and stromal populations. (**E)** Distribution of immune populations across selected MM1 and MM2 domains. (**F)** Distribution of stromal populations across selected MM1 and MM2 domains.

### Osteolytic and profibrotic processes are exclusively associated with the del(17p) MM1 clone

The MM1 region displayed profound remodelling of the bone marrow niche characterised by extensive osteolytic destruction and fibrosis **(Fig 5B)**. MM1 cells are exclusively located in the vicinity of osteolytic lesions, which is not observed with MM2 cells. Along with these histological changes, Xenium spatial transcriptomics demonstrated a marked enrichment of stromal and bone-associated populations within the MM1 region. Osteoblasts were increased approximately sixty-fold compared with region 2 **(Fig 5C)**, while osteoclasts, mesenchymal stromal cells (MSC/CAR and MSC/TAF), tumour-associated endothelial cells and perivascular endothelial cells were also substantially enriched. In contrast, the MM2 region contained relatively few stromal elements and was enriched for a population of resident macrophages (TAM2), suggesting preservation of a more homeostatic marrow environment **(Figs. 5C-F).** The MM1 region was characterised by enrichment of ECM and matrix-remodelling proteins including OGN, COL1A1, COL2A2, FN1, TGFB1, SERPINH1, FBLN1, LTBP2, FRZB and EMILIN1 **(SI File 3)**, consistent with a collagen-rich, TGF-β-associated stromal niche. This profibrotic, osteolytic and angiogenic region spatially co-localised with a TAM1-enriched macrophage compartment **(Fig. 5E)**. In contrast, the MM2 region was enriched for plasma cell-associated and immunomodulatory proteins, including SDC1, CD38, SLAMF1, ZBP1, KMO and ADA2 **(SI File 3)**, together with oxidative phosphorylation **(Figs. 4C-E)**, consistent with a metabolically active tumour compartment associated with a TAM2 macrophage niche.

The osteoblastic compartment in the MM1 region exhibited a transcriptional programme consistent with activated matrix-remodelling cells rather than mature bone-forming osteoblasts. These cells expressed TWIST1, WNT5A, SP7, RUNX2, COL5A2, FN1, LUM, DCN, ACTA2 and TNFSF11 (RANKL), together defining a pro-fibrotic, osteogenic stromal phenotype associated with extracellular matrix deposition and osteoclast activation **(Fig. 5D and SI Fig.7B)**. Expression of PDGFRA, PDGFRB, KITLG, IL1R1 and CSF1 further suggested extensive bidirectional communication with myeloid populations and osteoclast precursors (**Fig. 5D and SI Fig. 7B**). Complementing this programme, MSC/TAF cells demonstrated a classical activated fibroblast signature including VCAN, TNC, FBN1, C1R, C1S, FN1, DCN, THY1, PDGFRA, PDGFRB and ACTA2 **(Fig 5D, SI Fig 7B)**, indicating establishment of a fibrotic stromal niche capable of supporting tumour persistence. The osteoclast compartment was similarly expanded within the MM1 region and displayed an activated bone-resorptive phenotype. High expression of TNFRSF11A (RANK), CSF1R, MMP9, PTGS1, SPI1, CCR1, VEGFA and LIF is consistent with enhanced osteoclast differentiation, extracellular matrix degradation and pathological bone resorption (**Fig. 5D and SI Fig. 7B**). Together with abundant stromal RANKL expression, these findings indicate local activation of the RANK/RANKL axis as a likely mechanism underlying the osteolytic changes observed histologically.

Endothelial populations were likewise expanded in the MM1 region and exhibited evidence of active vascular remodelling. Both tumour-associated endothelial cells and perivascular endothelial cells expressed angiogenic markers including VWF, FLT1, CLEC14A, PLVAP, CD34 and ENG, together with IL15, ENTPD1 (CD39), HLA-E and interferon-stimulated genes including ISG15, IFIT3, IFITM3 and MX1 (**SI Fig.7B**). These findings suggest establishment of an activated vascular niche with concurrent immune-regulatory properties capable of sustaining tumour growth while limiting effective lymphocyte function.

### Myeloma clones are associated with different immune interactions

Along marked stromal remodelling, the MM1 region was also enriched for lymphoid populations. CD8 T cells and NK-like cytotoxic T cells were increased by approximately seven-to ten-fold compared with region 2 **(Fig 5C)**, while CD4 T cells were also expanded. These lymphocytes expressed a tissue-resident activated cytotoxic programme characterised by CCL5, GZMA, GZMB, PRF1, FASLG, KLRK1, CXCR6, CD69 and EOMES, together with inhibitory receptors including PDCD1, TIGIT, LAG3 and TOX, consistent with chronic antigen stimulation and progressive functional exhaustion (**SI Fig. 7B**). The simultaneous expression of cytotoxic effector molecules and multiple checkpoint receptors suggests that these lymphocytes remain antigen experienced but are likely constrained by the local suppressive microenvironment.

The myeloid compartment differed substantially between the two niches. The MM2 region was relatively enriched for TAM2 macrophages, whereas MM1 contained a marked expansion of TAM1 macrophages (**Fig. 5E**). Although both populations shared canonical macrophage markers, they exhibited distinct transcriptional identities. TAM1 cells expressed TREM2, CLEC10A, MSR1, FCGR2A, FCGR2B, VSIG4, GPR34, CSF1R, CD163, LGALS9 and PDCD1LG2 (**SI Fig. 7B**), which were consistent with a lipid-associated immunoregulatory macrophage phenotype which has recently been implicated in tumour progression across multiple solid cancers and haematological malignancies^26^. In contrast, TAM2 cells retained features of inflammatory resident macrophages, expressing CXCL9, CXCL10, VCAM1, CD86, CD274, IRF5 and IRF8, suggesting greater antigen-presenting capacity despite concurrent expression of several suppressive markers. The preferential accumulation of TREM2-positive macrophages within the del(17p) niche suggests selective recruitment or differentiation of highly suppressive macrophages^27^ by the aggressive clone. An unexpected feature of both tumour regions was the near-complete exclusion of granulocytes despite their abundance elsewhere within the trephine **(Fig. 3G and SI Fig. 6E)**, an observation previously noted in another study^28^. This indicates that both myeloma clones establish sharply defined immune boundaries, although the biological mechanisms appear distinct. The del(17p) clone in the MM1 region is associated with profound stromal activation, angiogenesis, osteoclastogenesis and accumulation of suppressive TREM2-positive macrophages despite abundant infiltrating cytotoxic lymphocytes, whereas the gain(12q) clone in the MM2 region occupies a comparatively quiescent marrow niche with fewer stromal alterations and predominance of resident macrophages. Taken together, the data are consistent with a myeloma microenvironment containing a fibrotic stromal niche (MSC/TAF) that is likely driving extracellular matrix remodelling and RANKL-mediated osteoclastogenesis, alongside immunosuppressive TAM1 macrophages that reinforce osteoclast activation and matrix remodelling. This combination is characteristic of aggressive myeloma-associated bone disease, where activated fibroblast-like stromal cells and M2-like macrophages cooperate to promote both fibrosis and osteolytic bone destruction.

## DISCUSSION

This study provides a proof-of-concept for integrated, multi-modal spatial profiling of multiple myeloma using a single diagnostic bone marrow trephine. We demonstrate that genomic, transcriptomic, proteomic and single-cell analyses can be combined to generate a coherent, high-resolution view of tumour architecture and its microenvironment. While previous studies have independently applied spatial transcriptomics, single-cell sequencing, bulk proteomic or genomic profiling to myeloma^14, 15, 17, 24, 29–35^, our work illustrates the added value of integrating these complementary modalities on the same clinical specimen. Spatial whole-genome sequencing defined clonal architecture, spatial transcriptomics linked genetic clones to distinct tumour, stromal and immune cell states, while single-cell RNA sequencing and immune profiling independently validated cellular composition and transcriptional programmes. Spatial proteomics uniquely captured extracellular matrix organisation and tissue remodelling incompletely represented at the transcript level. Together, this framework provides a comprehensive view of the genomic, cellular and microenvironmental interactions underlying spatial heterogeneity in multiple myeloma, revealing relationships obscured by any single technology. We show that myeloma subclones occupy distinct, spatially organised niches with unique molecular and microenvironmental features, supporting a model in which tumour evolution occurs within structured ecosystems rather than randomly.

A principal finding is the spatial coupling between genomic heterogeneity and transcriptional state. Shallow whole-genome sequencing identified two major tumour clones, including a clone harbouring del(17p) (MM1) and a genetically distinct clone characterised by gain of chromosome 12q (MM2). These genomic populations corresponded closely with transcriptionally distinct plasma cell states identified by spatial transcriptomics, suggesting that genetic evolution is accompanied by local functional diversification within the marrow niche. This link extended to individual genes on the altered chromosome arms. The del(17p) clone showed reduced TP53, whereas the gain(12q) clone showed increased MDM2, the main negative regulator of p53. Both clones may therefore weaken p53 signalling through different genomic routes. This concordance supports a role for spatially constrained microenvironmental interactions in shaping the phenotypic consequences of genomic evolution. Previous genomic studies have established extensive branching evolution in myeloma^36, 37^, while spatial transcriptomic studies have demonstrated regional transcriptional diversity within myeloma lesions^32, 33^. Our findings extend these observations by directly linking spatial genomic clones with corresponding transcriptional phenotypes within intact tissue.

Recent single-cell and spatial studies have shifted the view of multiple myeloma from a genetically heterogeneous tumour within a passive bone marrow niche to a dynamic ecosystem characterised by reciprocal interactions between malignant plasma cells and stromal, immune and osteolineage compartments^15, 30^. Stromal contact can remodel the chromatin landscape and transcriptional state of myeloma cells^29^, while inflammatory mesenchymal populations establish signalling niches that support tumour persistence^30^. Our integrated spatial proteomic and transcriptomic analyses extend this model by suggesting that genetically distinct myeloma clones are associated with discrete extracellular matrix environments. Specifically, in our case the del(17p) clone localised to a collagen-rich, osteogenic matrix-remodelling niche, whereas the gain(12q) clone occupied a transcriptionally and proteomically distinct stromal compartment. These findings support an ecological model in which tumour clones and their surrounding microenvironments undergo reciprocal adaptation, generating spatially segregated tumour ecosystems within the same bone marrow. Our data thus reinforces the recent observation of myeloma evolution capable to develop a spatially defined microenvironment^38^.

The immune microenvironment similarly exhibited marked spatial organisation. Despite abundant immune infiltration, transcriptional profiling suggested an immunologically “hot” yet functionally suppressed environment. Myeloid populations appeared central to this phenotype, with macrophages and granulocytes occupying regions enriched for immunosuppressive signalling. In particular, macrophages associated with the del(17p) clone displayed features consistent with alternatively activated tumour-associated macrophages, together with programmes linked to fibrosis and osteolytic remodelling. These observations agree with growing evidence that myeloid cells are major regulators of immune evasion, angiogenesis and bone destruction in MM rather than passive bone marrow resident components^31, 39, 40^.

The cytotoxic lymphoid compartment demonstrated additional mechanisms of immune dysfunction. Rather than classical checkpoint signatures alone, tumour regions exhibited altered expression of non-classical MHC-I molecules, including HLA-E and HLA-F, together with stress-induced ligands including MICA, MICB, ULBP family members and RAET proteins. These pathways may modulate natural killer cell and CD8⁺ T-cell activity through interactions with inhibitory receptors such as NKG2A and KIRs, suggesting mechanisms of immune escape not fully captured by conventional checkpoint analyses. Such findings are consistent with increasing recognition that non-classical antigen presentation pathways contribute to resistance against immune surveillance in haematological malignancies^41^.

More broadly, our work provides a framework for studying how genetic diversity and tissue architecture interact, with potential to guide therapeutic approaches by identifying spatially defined tumour vulnerabilities. The multiple immunosuppressive pathways identified in our example, including BTLA, MIF, MDK, LAIR1, TREM2, CD39/CD73, HLA-E and HLA-F, underscore the extensive immune regulatory landscape of the bone marrow microenvironment. These pathways act through complementary mechanisms to impair cytotoxic lymphocyte function, promote immunosuppressive myeloid populations and facilitate immune escape, suggesting that immune dysfunction in multiple myeloma is sustained by redundant suppressive networks. Therapeutic strategies targeting these axes are gaining momentum, with BTLA^42^, TREM2^27^, LAIR1^43^ blockade advancing through early-phase clinical evaluation, while MIF or adenosine pathway (CD39, CD73)^44^ and MDK^45^ inhibitors have shown promising immunomodulatory activity in preclinical models. In parallel, inhibition of the HLA-E/NKG2A axis has shown the capacity to restore NK-and CD8⁺ T-cell activity and is being investigated in combination with established immunotherapies^46^. Although HLA-F targeting remains at an earlier stage, the concurrent expression of these immune checkpoints supports combinatorial approaches integrating novel immune modulators with CAR T-cell therapies and bispecific antibodies to overcome immune resistance and improve response durability in multiple myeloma.

A notable strength of this study is the incorporation of spatial proteomics. While transcriptomic measurements capture cellular programmes, proteins provide a more direct readout of tissue structure and function, particularly within the extracellular matrix. Spatial proteomics identified distinct ECM-rich regions concordant with transcriptionally and genomically defined niches while additionally revealing matrix remodelling incompletely reflected at the RNA level. This highlights an important advantage of integrated proteogenomic approaches, particularly in tissues such as bone marrow where matrix composition profoundly influences plasma cell survival, drug resistance and immune cell trafficking^12, 13, 15, 29, 30^. Future studies incorporating phosphoproteomics or metabolomics may further refine understanding of functional signalling networks within these spatial niches.

The principal novelty of this work lies not in any individual technology, but in integrating multiple orthogonal measurements from the same diagnostic specimen. Previous landmark studies have separately characterised MM using single-cell transcriptomics^24, 35, 47^, spatial transcriptomics^15, 29, 30, 32, 33^ or genomic evolution^36, 37, 48^. However, to our knowledge, no studies have directly aligned spatial genomics, high-plex spatial transcriptomics, proteomics and matched single-cell immune profiling within an individual myeloma patient. This demonstrates that cross-validation across modalities increases biological confidence while allowing each platform to compensate for the limitations of the others.

Several limitations should nevertheless be acknowledged. As a proof-of-concept investigation, conclusions are derived from a single patient and biopsy, and cannot capture the full biological diversity of myeloma. Larger clinically annotated cohorts will be essential to determine the reproducibility of identified spatial programmes and establish associations with molecular subtype, treatment response and clinical outcome. Although the custom 480 and 5100 marker gene Xenium panels enabled focused interrogation of disease-relevant pathways, targeted panels inevitably restrict discovery compared with whole-transcriptome approaches. Larger cohorts, serial treatment sampling and integration with longitudinal clinical outcomes will be particularly valuable for understanding temporal evolution of spatial ecosystems and mechanisms of therapeutic resistance.

While the chromosome 17 deletion in our sample was concordantly identified, neither spatial whole-genome sequencing nor sc-RNA-seq-derived single-cell copy-number analysis of the paired sample detected the reported 1q gain identified by diagnostic FISH (∼20% positive cells). This discrepancy may reflect spatial intratumour heterogeneity, with the 1q gain confined to a geographically restricted subclone captured by targeted FISH of the contemporaneous liquid aspirate but absent from the regions analysed by sequencing of the trephine specimen. Such spatial segregation of genetically distinct myeloma subclones has been described previously^37, 48^ and highlights the impact of sampling on genomic profiling.

Looking forward, integrated spatial multi-omics has considerable potential to transform biological discovery and clinical diagnostics^49–51^. Rather than classifying disease solely according to recurrent genomic abnormalities, future patient stratification may incorporate spatially resolved ecological features, including clonal organisation, immune suppression, stromal remodelling and matrix composition. Such multidimensional biomarkers may better predict therapeutic response than individual molecular features alone. Combining orthogonal spatial and single-cell technologies could uncover novel tumour–microenvironment interactions, identify spatially restricted biomarkers, and inform more effective, context-aware therapeutic strategies in myeloma and other hematologic malignancies.

However, translation into routine diagnostics will require simplified workflows, robust analytical pipelines and prospective demonstration of clinical utility and cost effectiveness. Artificial intelligence and machine learning will likely be instrumental by integrating heterogeneous imaging and molecular datasets, identifying latent spatial features beyond human interpretation and generating clinically actionable predictions^52^. Foundation models trained across histology, spatial transcriptomics and molecular pathology may ultimately infer complex molecular states directly from routine diagnostic sections, reducing costs while preserving biological resolution. As spatial technologies become increasingly scalable, integrated multi-modal profiling offers a promising framework for redefining multiple myeloma as a spatially organised ecosystem in which tumour evolution, immune dysfunction and stromal remodelling are intrinsically interconnected.

## MATERIAL AND METHODS

Studies were conducted in accordance with the Declaration of Helsinki and Good Clinical Practice guidelines, and were approved by the relevant national, regional and institutional review boards. The patient provided written informed consent for use of their samples. Ethics committees overseeing the Oxford Radcliffe Biobank and HaemBio banks approved sample collection/use. Bone marrow samples were acquired as part of routine care, including histopatholgical analysis and were stored within the Oxford Radcliffe Biobank (REC reference 19/SC/0173) and HaemBio (REC reference 24/EM/0060) with patient consent for future research.

### Sample processing

Whole bone marrow aspirate collected from a relapsed myeloma patient (Supplementary File 1) was passed through a CD138 bead selection kit (EasySep, StemCell). The flow-through (CD138-fraction) was separated on a Ficoll gradient, and the buffy coat was isolated, counted, and cells were viably frozen in DMSO/foetal calf serum (10/90, v/v). The cells retained on the beads were eluted, counted and ∼ 500,000 were pelleted in buffer RLT (Qiagen) viably frozen and stored in liquid nitrogen until use. Vials with BM aspirates were defrosted in small batches in a 37°C water bath for approximately 1 minute, then placed on ice and transferred dropwise into 4°C RPMI 1640 medium supplemented with 1% glutamine (Gibco) (concentration), and 10% FCS (v/v). Samples were washed twice by centrifugation at 300 × g for 5 minutes. Cells were resuspended in sterile PBS up to a concentration of 1 million per mL and passed through 70 µm filter tips (Bel-Art Flowmi).

Samples were immediately proceeded to parallel mass cytometry staining and single-cell transcriptomics workflows. A bone marrow trephine biopsy was collected from the same patient as part of routine clinical procedures by trained clinical staff according to standard NHS workflows. The trephine core was fixed, formic acid decalcified and embedded in paraffin to generate formalin-fixed paraffin-embedded (FFPE) bone marrow trephine blocks. Sections (5 µm-thick) were cut for the following spatial transcriptomics, LCM aided spatial proteomics and shallow whole genome sequencing.

### In situ imaging spatial transcriptomics

BM trephine sections were assessed by 480-gene and 5.1 K custom Xenium panels (**SI File 1**). Slides analysed with the 480-gene panel (custom designed myeloma immuno-oncology panel) were processed with Xenium v1 chemistry and reagents (10x Genomics). Slides analysed with the 5.1K-gene panel were processed with Xenium Prime chemistry and reagents (10x Genomics). The predesigned Xenium Prime 5K Human Pan Tissue & Pathways Panel (PN-1000724, 10x Genomics), targeting 5,001 genes, was supplemented with a custom add-on panel comprising 100 additional immuno-oncology genes. Multimodal cell segmentation staining was performed for both the 480-and 5.1K-gene panels (PN-1000661, 10x Genomics).

### Histology

Sections selected for spatial proteomics and genomics were cut on to PEN membrane slides (Thermofisher). The sections was deparaffinized in Clearene for 4 minutes, followed by rehydration through a graded ethanol series (100% twice, 80%, and 70%, each for 2 minutes) and distilled water (2 minutes). Sections were stained with Mayer’s Haematoxylin for 2.5 minutes, rinsed twice in distilled water, and treated with Bluing Reagent for 10–15 seconds before further rinsing. After a brief dip in 100% ethanol, sections were counterstained with Eosin for 2–3 minutes. Slides were subsequently washed in 100% ethanol (brief rinse), 80% ethanol (2 minutes), 70% ethanol (2 minutes), and water (2 minutes). For imaging, sections were temporarily mounted with 50% glycerol in DEPC water and scanned at 80x magnification using MoticEasyScan (Motic). Following scanning, coverslips were gently removed by immersion in water for 5–10 minutes, and the sections proceeded to dehydration through an ethanol series (70/80/100 %), followed by 20 minutes of air drying.

### Topological alignment of trephine sections

Digital HE images were exported in QuPath^53^, and digital laser cutting grids (coordinates of cutting areas) of equal size mini-tiles (250µm-wide) with 25µm interval were made. Images were aligned using the thin plate spline algorithm (TPS)^20^ with 8 manually selected calibration points via scikit-image library (v0.24.0)^54^ in Python. For integration analysis with xenium, coordinates of digital cutting grids were identified using TPS. Four adjacent mini tiles were pooled representing a 560µm tile area unit. Geojson files of the laser cutting grid were then transformed into xml format via py-lmd package (https://pypi.org/project/py-lmd/) in Python with added well id information indicating the match between tiles and plate wells. XML cutting outlines were imported into the Leica LMD Software (v8.x), controlling a Leica LMD7 system for automated laser microdissection.

### LCM enabled spatial whole genome sequencing

Microdissected tissue from spatial tile regions collected at the bottom of 96-well plates was briefly centrifuged (1,500 × g, 20 s), vortexed, and centrifuged again (1,000 × g, 10 s) to localize captured tiles to the bottom of the wells. Samples were treated by adding 10 µL per well of custom lysis buffer (30 mM Tris HCl pH 8.0, 0.5% Tween 20, 0.5% NP40, 25 µg/ml Proteinase K) followed by incubation in a thermal cycler (Thermo Fisher Scientific) with at 55°C for 15 min and 75°C for 15 min to promote lysis. Following this, the samples were dried by removing the plate seal and incubating on a thermal cycler at 75°C for 30 min with the lid open. After drying, genomic DNA was fragmented, end-repaired, and dA-tailed directly in the wells using NEBNext Ultra II FS DNA Module (NEB, Cat# E7810S/L). Separately annealed adapters were ligated in situ using the NEBNext Ultra II Ligation Module (NEB, Cat# E7595S/L). Ligated products from each 96-well plate were pooled and purified using AMPure XP beads (Beckman Coulter, Cat# A63880). The pooled ligation product was USER-treated (NEB, Cat# M5505S/L) and PCR amplified for 20 cycles using NEBNext Ultra II Q5 Master Mix (NEB, Cat# M0544S/L) with i5 and i7 indexing primers.

Amplified libraries were purified by bead cleanup and assessed using High Sensitivity D1000 ScreenTape (Agilent Technologies, Cat# 5067-5584) with High Sensitivity D1000 Reagents (Agilent Technologies, Cat# 5067-5585) on an Agilent TapeStation system. Where required to remove residual adapter dimers, libraries were subjected to 3% agarose gel purification using GelRed Agarose LE (Biotium, Cat# 41029-5G), and DNA fragments <200 bp were excluded. Size-selected DNA fragments >200 bp were excised and recovered using the Zymoclean Gel DNA Recovery Kit (Zymo Research, Cat# D4002). Purified multiplexed libraries were sequenced on an Illumina NextSeq 500 platform using a NextSeq 500/550 Mid Output Kit v2.5 (Illumina, Cat# 20024904). Raw demultiplexed reads were demultiplexed further using Cutadapt (v4.4) to obtain well-level fastq files which were quality-trimmed and aligned to the reference genome using BWA-MEM. Alignments were coordinate-sorted and deduplicated using SAMtools to generate clean BAM files for downstream CNV analysis.

### LCM enabled spatial LC-MS/MS proteomics

Micro-dissected tissues of were transferred into wells of a 96-well PCR plate (Eppendorf, cat. no. 0030129580). To collect tissue at the bottom of the well, 150 µL acetonitrile was added to each well, the plate sealed and centrifuged at 2000 *g* for 5-10 minutes, dried in a vacuum centrifuge at 40 °C, and stored at - 20 °C until further processing. Tissue areas were lysed in 20 µL of lysis buffer 0.013% n-Dodecyl-β-D-Maltoside (DDM, Thermo Fisher Scientific, cat. No. 89902), 50 mM triethylammonium bicarbonate (TEAB, Sigma, cat. No. 18597) in LC-MS/MS-grade water and incubated at 95 °C for 90 minutes. Samples were cooled to 20 °C and centrifuged at 2000 *g* for 1 minute. Five µL of 60% acetonitrile in 50 mM TEAB was added to each sample and incubated at 75 °C for 30 minutes, cooled to 20 °C and centrifuged at 2000 *g* for 1 minute. A further 4 µL of 50 mM TEAB was added to dilute the acetonitrile to 10%, then 1 µL of 8 ng/µL Trypsin/LysC enzyme mix (Promega, cat. No. V5073) in 50 mM TEAB was added to the samples. The plate was sealed and incubated at 37 °C for 16 hours in a thermal cycler with the lid set to 50 °C. The digestion was stopped by adding formic acid to 1 % (v/v), and the plate stored at −20 °C until analysis. Before analysis by LC-MS, samples were diluted by adding 5 μL of sample to 15 μL solvent A (0.1 % formic acid in water). Diluted samples were loaded on Evotip Pure C18 tips (Evosep, EV2011) as per manufacturer’s protocol. Briefly, tips were rinsed with 20 µL 99.9 % acetonitrile 0.1 % formic acid by centrifugation, conditioned by soaking in 1-propanol, equilibrated with 20 µL Solvent A by centrifugation, loaded with 3 µL of diluted sample and 17 µL of solvent A, then centrifuged, washed with 20 µL solvent A by centrifugation, and wetted with 100 µL solvent A and a brief centrifugation step for 10 seconds. All centrifugation steps were 60 seconds at 800 *g*. Peptides were analysed on an Evosep One LC system (EvoSep) coupled to a timsTOF Ultra 2 mass spectrometer (Bruker) using the Whisper Zoom 40 samples-per-day method and a 75 µm × 150 mm C18 column with 1.7 µm particles and an integrated Captive Spray Emitter (IonOpticks). Buffer A was 0.1% formic acid in water and Buffer B was 0.1% formic acid in acetonitrile. Data were collected using diaPASEF with 1 MS frame and 8 diaPASEF frames per cycle with an accumulation and ramp time of 100 ms, for a total cycle time of 0.96 seconds. The high sensitivity parameter was enabled. The diaPASEF ^55^ frames were separated into 3 ion mobility windows, in total covering the 400–1000 m/z mass range with 25 m/z-wide windows between an ion mobility range of 0.64–1.45 Vs/cm^2^. The collision energy was ramped linearly over the ion mobility range, with 20 eV applied at 0.6 Vs/cm^2^ to 59 eV at 1.6 Vs/cm^2^. Mass spectrometry raw files were analysed in DIA-NN version 1.9.2 using an in-silico spectral library generated by DIA-NN with default settings, except that carbamidomethylation of cysteine was not selected, allowing for 1 missed cleavage. A Uniprot human FASTA file containing 20,514 sequences, with common contaminants added by DIA-NN^56^ was used. Mass accuracy for MS1 and MS2, and scan window were set to 0 (automatic determination), the FDR threshold was set to 1 %. Protein intensities from DIA-NN’s ‘pg_matrix’ output were used in the further downstream analysis.

### Long reads single cell transcriptomics

Bone marrow aspirate cells were processed as described^57^. In brief, cells were treated using GEM-X beads (10X Genomics), with cDNA library amplification per manufacturer’s protocol (10X Genomics, user guide CG000731). cDNA libraries were quantified with TapeStation and 10 ng cDNA taken forward into the Oxford Nanopore (ONT) protocol using PCS114 chemistry on a Promethion P24 using R10.4.1 flow cells (ONT). Super-high accuracy basecalling was performed by Dorado within the MinKNOW software.

### Immune mass cytometry (IMC) imaging

FFPE sections (5 mm) were cut onto glass slides and baked overnight at 60°C prior to deparaffinisation in xylene for 20 minutes. Slides were then rehydrated through a graded series of ethanol solutions for 5 minutes in 100%, 95%, 80% and 70% EtOH. Slides were then washed twice in Maxpar water for 5 minutes each prior to antigen retrieval. Antigen retrieval was performed by submerging the tissue section in pH 9.0 antigen retrieval solution (Ebiosciences) diluted to a 1x working solution in Maxpar water. Slides were then incubated at 95°C for 30 minutes in a thermocycler and allowed to cool to RT before washing twice in Maxpar PBS (MPPBS) for 5 minutes each. Sections were then blocked with SuperBlock blocking solution (Thermo scientific) for 30 minutes at RT. Tissues were then stained with metal-tagged antibody master mix overnight at 4°C in a hydration chamber. Sections were then washed in 0.1% Triton-X (sigma) diluted in MPPBS twice for 8 minutes each followed by two further 8-minute washes in MPPBS. Sections were then stained with Cell-ID Ir-Intercalator (Standard BioTools) diluted 1:500 in MPPBS for 30 minutes at RT. The tissue sections were then washed with deionised water for 5 minutes then air dried at RT. Regions were acquired using the Hyperion Imaging System (Standard BioTools) with pulsed laser ablation performed at 200 Hz. Image analysis was performed using the HALO suite (Indica Labs).

### Mass cytometry

Up to 3 million cells per sample were washed twice in Maxpar Cell Staining Buffer (CSB; Standard BioTools) with 5 minutes centrifugation at 300 × g and stained with pre-aliquoted surface and intracellular antibody cocktails (**Supplemental Table 5**). Prior to intracellular permeabilization, cells underwent an additional 1.6% formaldehyde fixation. Rhodium live/dead staining was included in the surface antibody staining reaction. Data were acquired on a Helios mass cytometer (Standard Biotools), according to the manufacturer’s protocol. Normalized .fcs files were uploaded into Cytobank flow cytometry software and processed as previously described ^58^. Singlet events were exported as .fcs files and processed in R (version 4.4.1, R Core Team 2024). QC and visualization was performed with CATALYST; batch correction with CyCorrect and differential abundance analysis with MiloR.

## Data Analysis

### Spatial transcriptomics

The Xenium 5.1K and 480 marker datasets were analysed using Seurat (v5)^59^. Cell-feature matrices together with the corresponding centroid and segmentation coordinates were imported to generate a Seurat object for quality control, clustering and manual cell-type annotation. Cell segmentation was refined using FastReseg^60^, with integrated single-cell sequencing (scRNA-seq) data^57^ as the reference for both resegmentation and automated cell-type annotation. The Seurat object was subsequently reconstructed using the updated polygons generated with cellPoly (github.com/Nanostring-Biostats/CosMx-Analysis-Scratch-Space/tree/Main/_code/cellPoly) and the updated cell-feature matrix. Following initial annotation, major cell clusters were further refined by sub-setting, re-clustering, and manual annotation based on canonical marker gene expression.

Neighbourhood analysis was performed using BANKSY^61^ and Squidpy (v1.8.3)^62^ in combination with the FindNeighbors function implemented in Seurat to identify spatial domains, cellular niches, and the localisation of myeloma cell populations. Developmental trajectories of cell populations were inferred using Slingshot^63^. Cell-cell communication analysis was performed using CellChat (v2)^64^. For ligand-receptor interaction inference, the maximum interaction distance was set to 250 mm, with contact-dependent signalling defined within 10 mm and secreted signalling within 100 mm. Differential expression analysis among cell clusters was performed using the FindAllMarkers function within the Seurat package in R, and cluster-specific marker profiles were displayed using the DotPlot function. For high-resolution spatial visualization, per-cell centroid coordinates alongside metadata (cell annotations and spatial domains) were extracted from the Xenium Seurat object and plotted in Python using Matplotlib^65^ with each point representing a cell at its physical centroid. To align spatial transcriptomics with adjacent laser capture microdissection (LCM) regions evaluated in downstream genomic and proteomic assays, tile-level pseudobulk RNA matrices were constructed by summing raw transcripts within each grid tile and normalizing by total tile cell count. Principal Component Analysis (PCA) and k-nearest neighbor (k-NN) clustering were performed on the normalized tile matrices, with k = 6 selected based on elbow plot inflection analysis. Cluster-specific markers identified via FindMarkers in Seurat, ranked by log2 fold-change and statistical significance, were then subjected to Gene Set Enrichment Analysis (GSEA) against the MSigDB Hallmark gene set using the fgsea^66^ package in R. Differential expression between the MM1 region tiles and MM2 region tiles was tested with DESeq2 on raw pseudobulk counts, using size factors derived from the total cell number of each tile.

### scRNAseq

Raw scRNA-seq FASTQ files were processed using SiCeLoRe^67^. Reads were aligned to the reference genome GRCh38 using minimap2^68^. Transcript-and gene-count matrices generated by SiCeLoRe workflow were used for downstream analysis in Seurat. Cells with >500 detected genes counts, >800 transcripts, ≤ 20% mitochondrial genes, and classified as singlets by scDblFinder^69^ were retained for downstream analyses. Data were normalised; dimensionality reduction using identified highly variable genes and clustering were performed using the standard workflow of Seurat. Cell-type annotation was performed manually based on expression of canonical marker genes. A copy number variation (CNV) profile was inferred using CopyKAT^70^.

### Spatial WGS

Aligned BAM files from all spatial tile samples were imported into the CopyKit R ^71^ package to calculate copy number profiles for each pooled spatial tile. Low-quality tiles failing quality control filtering were excluded from downstream analyses. The Louvain algorithm was applied to cluster spatial tiles based on their copy number profiles, identifying distinct genomic subclones that were visualized as CNV heatmaps. Phylogenetic relationships among identified subclones were manually reconstructed based on the evolutionary progression and shared segmental CNV features across spatial domains.

### Spatial proteomics

Raw protein intensity matrices from spatial tiles were imported into R. Tiles exhibiting a missing protein ratio exceeding 80% were filtered as low quality. The remaining expression matrix was log2-transformed, median-normalized, and imputed for low-abundance missing values via a left-shifted normal distribution using the DEP ^72^ package. Dimensionality reduction was performed by Principal Component Analysis (PCA), and tiles were partitioned by unsupervised k-means clustering (k = 6) on the first 30 principal components. k = 6 was selected based on the elbow plot together with inspection of the resulting spatial cluster patterns. Cluster-specific differential protein markers were identified using Wilcoxon rank-sum tests (one cluster versus all others) with Benjamini–Hochberg correction. Significant differential proteins (adjusted p < 0.05), ranked by log2 fold-change, were subjected to pre-ranked Gene Set Enrichment Analysis (GSEA) against MSigDB

Hallmark gene sets using the fgsea package. Parallel differential protein and GSEA pathway analyses were conducted across the corresponding spatial subclone clusters. To specifically evaluate extracellular matrix (ECM) dynamics, proteins were cross-referenced against the Matrisome database^25^. ECM-related proteomic profiles underwent k-means clustering and differential marker analysis, with top ECM markers displayed via dot plots.

### Data Integration

To integrate spatial multi-omic layers across aligned spatial tiles, a Multi-Omics Factor Analysis (MOFA) model was constructed in Python using the MOFA ^73^ framework. Input modalities comprised binned genomic CNV profiles (log2 ratio across 4107 genomic bins), pseudobulked spatial RNA matrices, and normalized proteomic intensity data. Trained model parameters and factor weight matrices for 15 latent MOFA factors were imported into R using the MOFA2 package. Unsupervised k-means clustering was performed on the factor weight matrix on tile level to identify integrated spatial multi-omic states. Variance decomposition was conducted to quantify the proportion of variance explained by each latent factor within the WGS, RNA, and proteomic modalities. Genes and proteins were ranked by their factor weights to identify the top contributors to each factor.

## Supporting information

https://github.com/jinsenlu-hub/Myeloma-project/tree/d0605d0bc140f78b473ec539f13aa86a16ef5bac/Supplementary%20file

## ACKNOWLEDGMENTS

Work in the UO laboratory is supported through research grants from BristolMyersSquibb, GlaxoSmithKline, the Oxford Translational Myeloma Centre, a program grant of the Bone Cancer Research Trust (BCRT11025) and the LEAN program grant of the Leducq Foundation. SR is funded by Prostate Cancer UK (RIA22-ST2-004). APC is a recipient of a Medical Research Council (MRC) career development fellowship (MR/V010182/1). AS is a recipient of JSPS Clinical PhD Fellowship, funded by the Jean Shanks Foundation and the Pathological Society. We acknowledge the contribution to this study made by the Oxford Centre for Histopathology Research and the Oxford Radcliffe Biobank, which are funded by the University of Oxford, the Oxford CRUK Cancer centre, and the NIHR South Central RRDN.

## Data availability

The LC-MS/MS proteomics data have been deposited to the ProteomeXchange Consortium via the PRIDE^74^ partner repository with the dataset identifier PXD083301; transcriptomic data are deposited with GEO (GSE307660).

All custom code and bioinformatic scripts generated for the analyses in this study are publicly available in the GitHub repository: https://github.com/jinsenlu-hub/Myeloma-project.git.

Standard bioinformatics tools and established software packages used for data processing are not hosted in this repository but are described in detail, along with their version numbers and parameters, within the Methods section.

## Author contributions

Conception: UO, SR, RF, JL. Supervision: UO, SR, RF. Experiments: JL, SD, EW, WB, EC, JH, PBB, SR. Materials, protocols: CJ, AS, JH, SG, SGr, FI, AL, CB, SuG, KR, AT, RC, DR, NA, SR. Data analysis: JL, CYW, EW, CJ, APC, RF, UO, SR. Funding and resources: UO, RF, SR. Manuscript draft: UO. All authors have read, approved and contributed to manuscript editing.

