## Supplementary material for "A platform for deep topographic multiomic mapping of the bone marrow environment in multiple myeloma": https://github.com/jinsenlu-hub/Myeloma-project/tree/d0605d0bc140f78b473ec539f13aa86a16ef5bac/Supplementary%20file

**SI File 1. Xenium in situ marker panels (480-gene and 5.1K).**
Sheet1. 480 (380+100 custom) -gene Xenium panel;
Sheet2. 5.1K (5,000 + 100 custom) gene Xenium panel;
Sheet3. 480-panel differential marker genes per cell cluster
**SI File 2. CNV analysis outputs from spatial low-coverage WGS of LCM tiles.**
Sheet1. Per-tile sequencing QC and metadata;
Sheet2. Raw read counts per genomic bin per tile;
Sheet3. Log2 R ratio (copy-number) per genomic bin per tile
**SI File 3. Total and matrisome protein expression from LC-MS of LCM tiles.**
Sheet1. Total protein expression matrix per tile;
Sheet2. Per-tile proteomics metadata;
Sheet3. Matrisome-matched proteins with annotation and expression
Sheet4. Protein differential analysis between MM1 and MM2 area.
**SI File 4. Mass cytometry (CyTOF) antibody panel for bone-marrow aspirate profiling.**
**SI File 5. Pseudobulk RNA expression and per-tile metadata from 480 Xenium assay.**
Sheet1. Pseudobulk RNA counts per tile;
Sheet2. Per-tile RNA cluster and cell-number metadata
**SI File 6. Geometry and cross-modality tile mapping for processed LCM tiles.**
**SI File 7. Genes and proteins with expression profile matches the subclone-specific copy number alterations of the MM1 and MM2 regions.**
Sheet1. CNV-concordant genes (RNA);
Sheet2. CNV-concordant proteins;
Sheet3. CNV-concordant proteins annotated to biological processes;
Sheet4. CNV-concordant genes annotated to biological processes

**SI Figure 1:** Histology and myeloma related clinical metadata.

**SI Figure 2:** CNV architecture of identified subclones derived from sWGS data.

**SI Figure 3:** Cell composition and annotation of single-cell RNA sequencing (scRNA-seq) data from bone marrow aspirate.

**SI Figure 4:** Cell composition and annotation of Xenium 480 dataset from bone marrow trephine.

**SI Figure 5:** Cell composition and annotation of Xenium 5.1K dataset from bone marrow trephine.

**SI Figure 6:** Granulocyte lineages identified in Xenium and single cell CyTOF datasets.

**SI Figure 7:** Immune-stromal marker expression of MM1/MM2 regions in Xenium 480 dataset.

**SI Figure 8.** Multi-Omics Factor Analysis (MOFA) integration of spatial WGS, proteomics and pseudobulk RNA data across LCM tiles.

**SI Figure 1: Histology and myeloma related clinical metadata.** **(A)** Representative H&E and immunofluorescence (IF) images of consecutive formalin-fixed paraffin-embedded (FFPE) sections. These sections were utilized for downstream whole-genome sequencing (WGS), liquid chromatography-mass spectrometry (LC-MS), and Xenium spatial transcriptomics. IF markers were selected to visualize the cell nucleus (DAPI), cell boundaries (ATP1A1, CD45, E-Cadherin), RNA (18S), and structural proteins (α-SMA, Vimentin). **(B)** Summary of the corresponding clinical metadata, including patient age, sex, multiple myeloma subtype, treatment history, and cytogenetic findings.

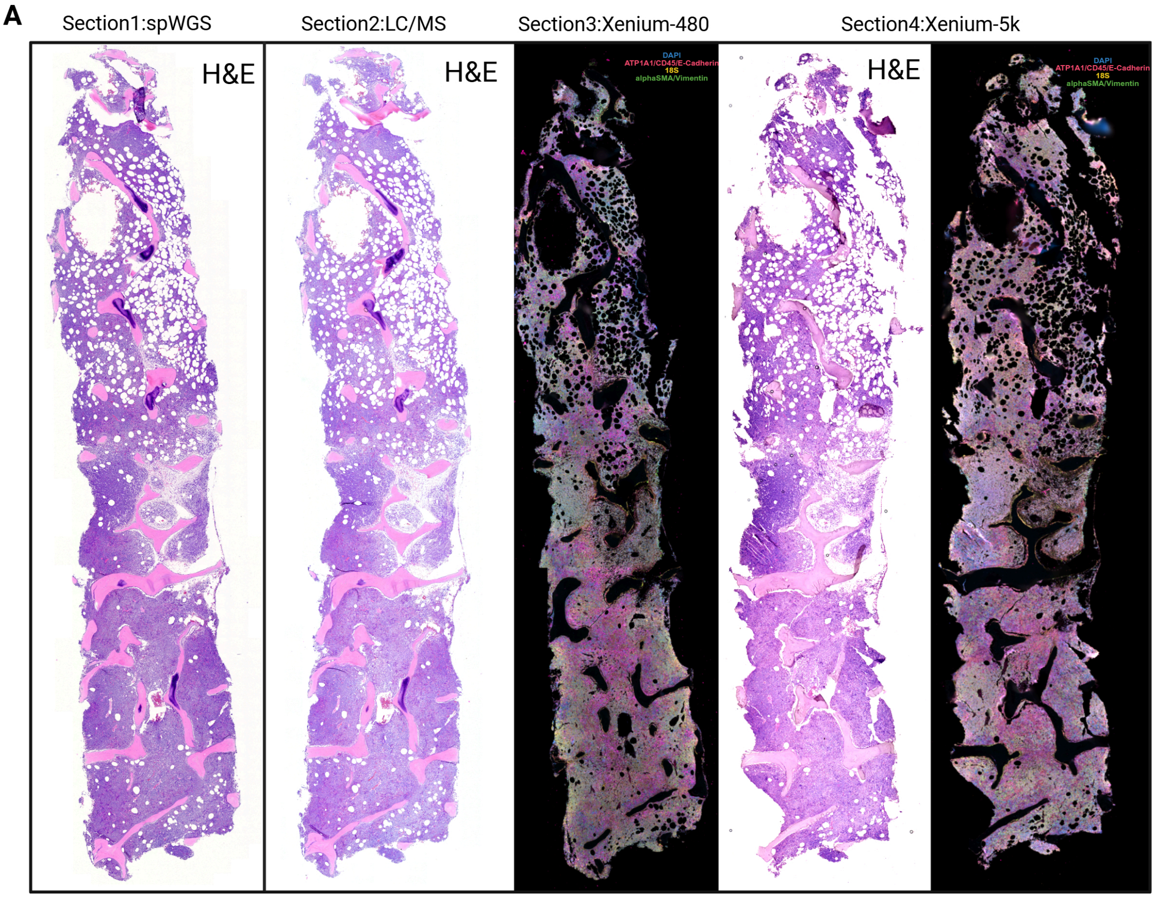


**B**

| | |
|---|---|
| **Age at treatment start (years)** | 59 |
| **Sex** | M |
| **Myeloma type** | Kappa |
| **Number of prior treatment lines** | 6 |
| **Prior treatment lines** | VCd_auto; IRd; D; Pomd; CC92480; DTPACE |
| **Cytogenetic findings** | gain 1q21 (22%); 14q32 rearrangement unknown partner (unknown %); del 17p13 (13%) |

**SI Figure 2. CNV architecture of identified subclones derived from sWGS data.** Consensus copy number profiles for five distinct cellular subclones (Subclones A–E). Across all panels, the x-axis represents chromosomal coordinates (chr1–22, X), and the y-axis represents the absolute total copy number. Gray dots depict the unsegmented, binned read counts, while the solid step-lines denote the algorithmically segmented consensus copy number states.

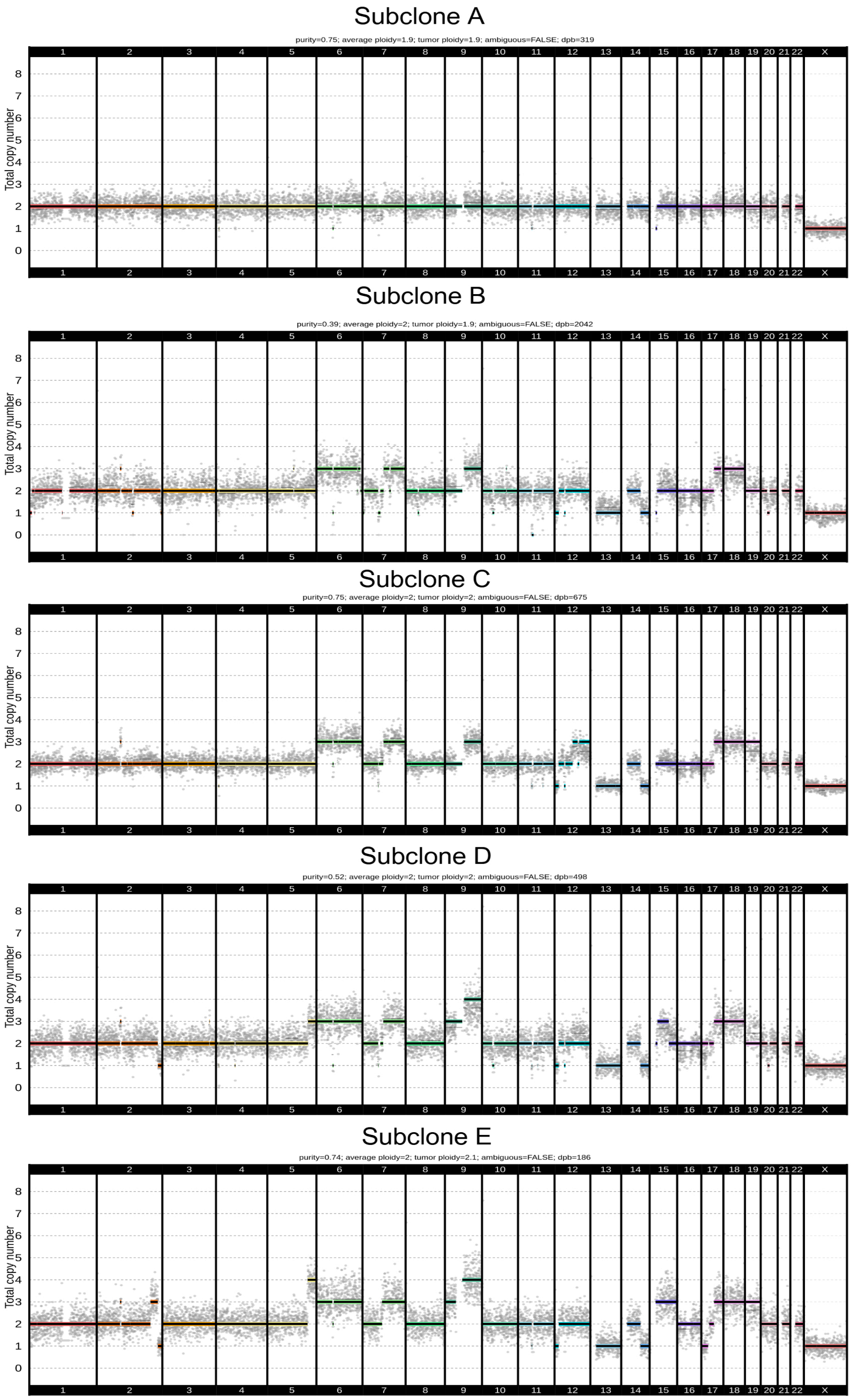

**SI Figure 3: Cell composition and annotation of single-cell RNA sequencing (scRNA-seq) data from bone marrow aspirate.** **(A)** UMAP plot showing the major cell populations identified in the bone marrow aspirate from the patient, including macrophages, neutrophils, B cells, T and NK cells. **(B)** Expression of canonical marker genes across each cell cluster. The relative proportions of each cell population is indicated on the right. **(C)** Heatmap showing the average expression of immune regulatory genes across the identified cell clusters.

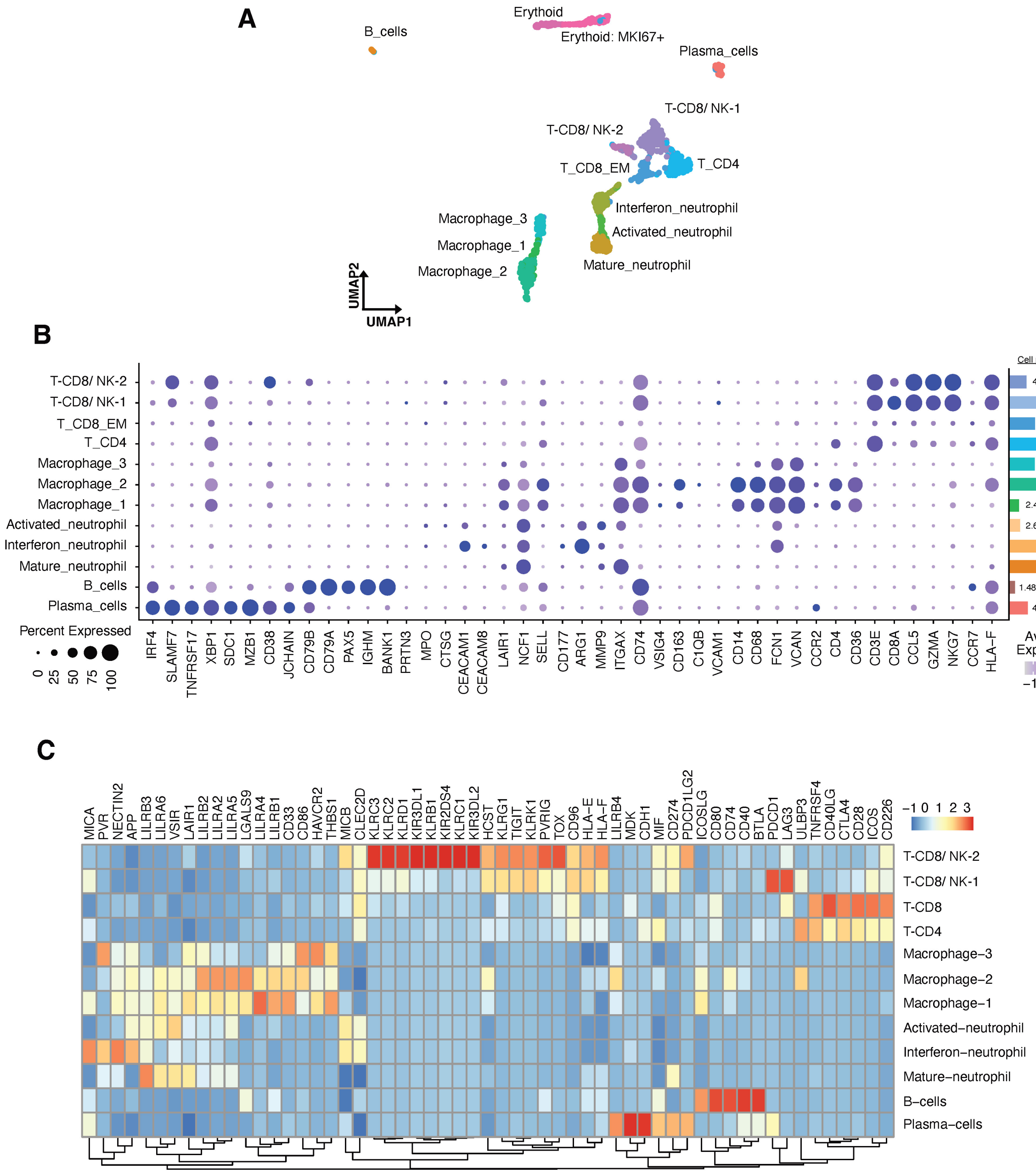

**SI Figure 4: Cell composition and annotation of Xenium 480 dataset from bone marrow trephine.** **(A-B)** UMAP of major cell types and specific immune **(A)** and stromal population **(B)** in the patient's bone marrow trephine. **(C-D)** The dot plot of gene expression profile of canonical markers **(C)** and top differential markers **(D)** for all identified cell types and niches with cell percentage illustrated as bar plot on the right side.

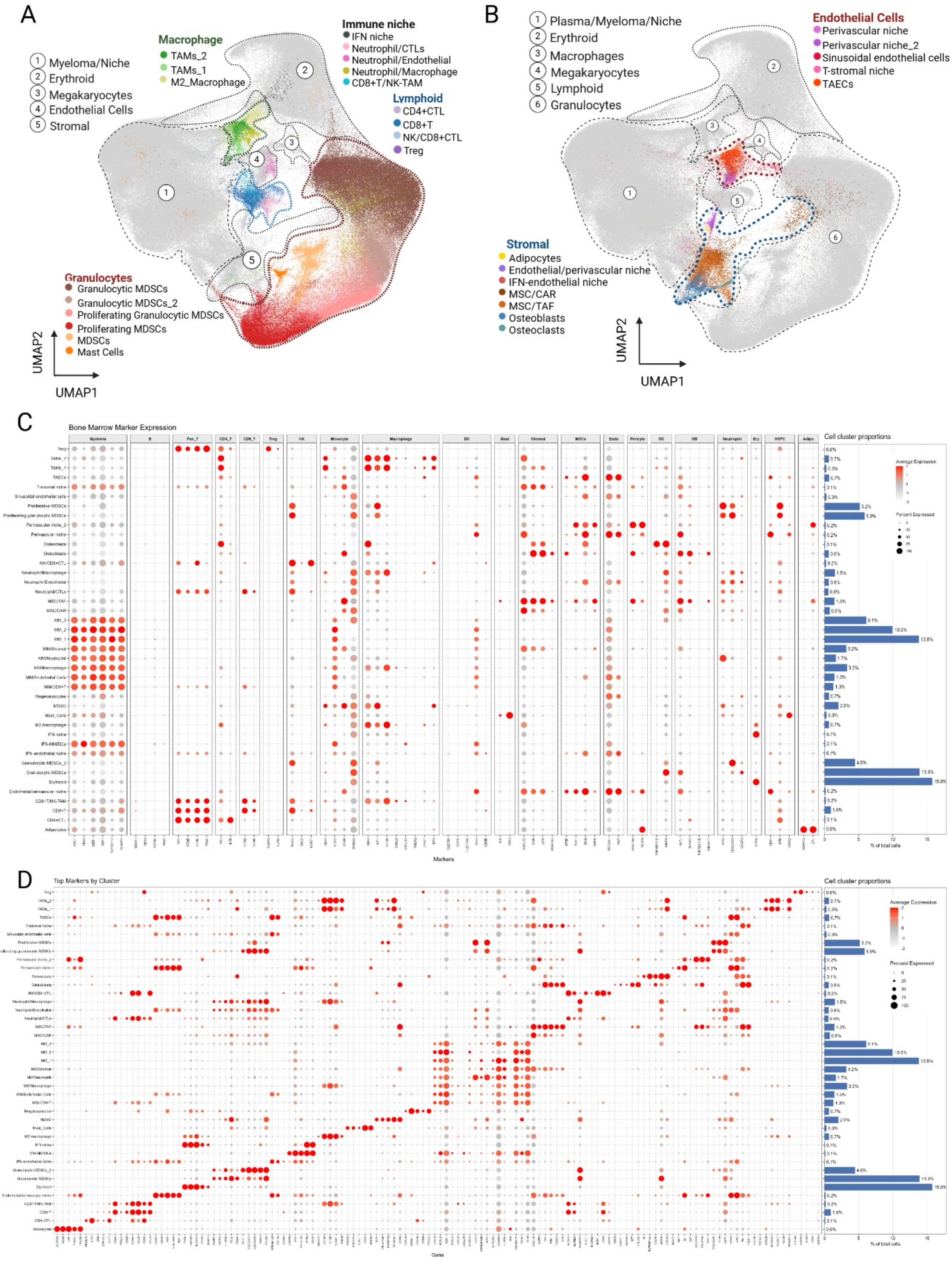

**SI Figure 5: Cell composition and annotation of Xenium 5.1K dataset from bone marrow trephine.** **(A)** UMAP plot showing the major cell types and their corresponding subtypes identified in the patient's bone marrow trephine. **(B)** Expression of canonical marker genes of the identified cell types across major cell cluster. The relative proportions of the identified cell types are indicated on the right.

A

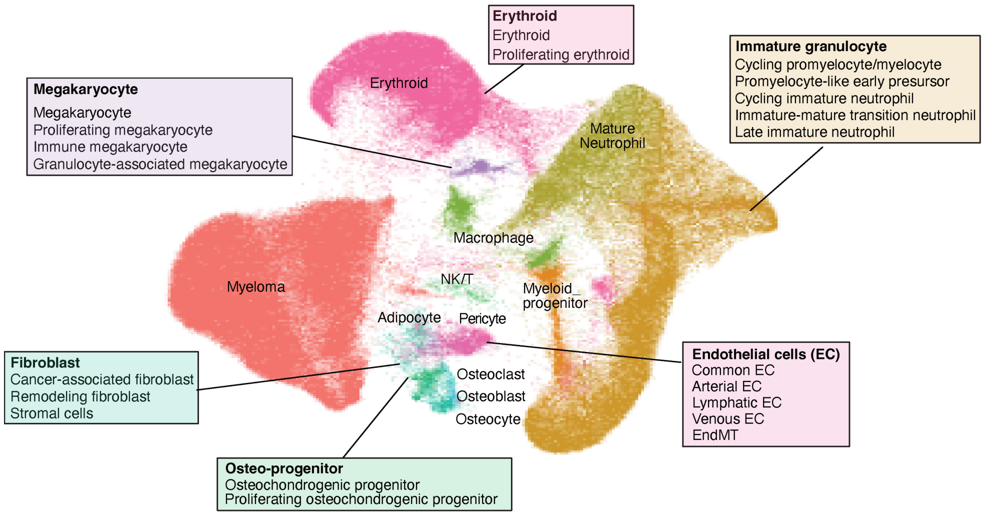


B

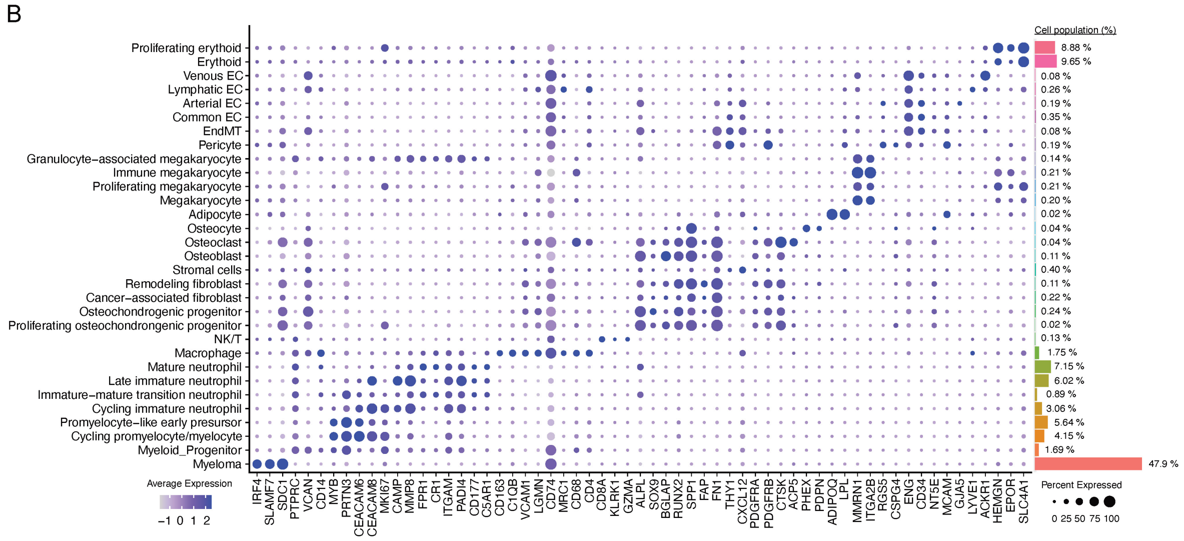

**SI Figure 6 Granulocyte lineages identified in Xenium and single cell CyTOF datasets. (A-B)** UMAP plots showing the granulocyte lineages identified in the Xenium 5K dataset **(A)** and their inferred pseudotime developmental trajectory **(B)**. **(C)** Heatmap showing the average expression of marker genes used to identify the granulocyte lineages. **(D)** Heatmap showing the average expression of immune-regulatory genes across myeloma and immune cells. **(E)** Spatial neighbourhood enrichment analysis between major myeloma cluster and granulocytes and selected immune population from xenium 480 dataset. **(F-G)** UMAP plot showing the granulocyte lineages identified by CyTOF **(F)** and heatmap showing the expression profiles of the CyTOF marker panel across granulocyte lineages, together with the relative proportions of each lineage within the neutrophil population **(G)**.

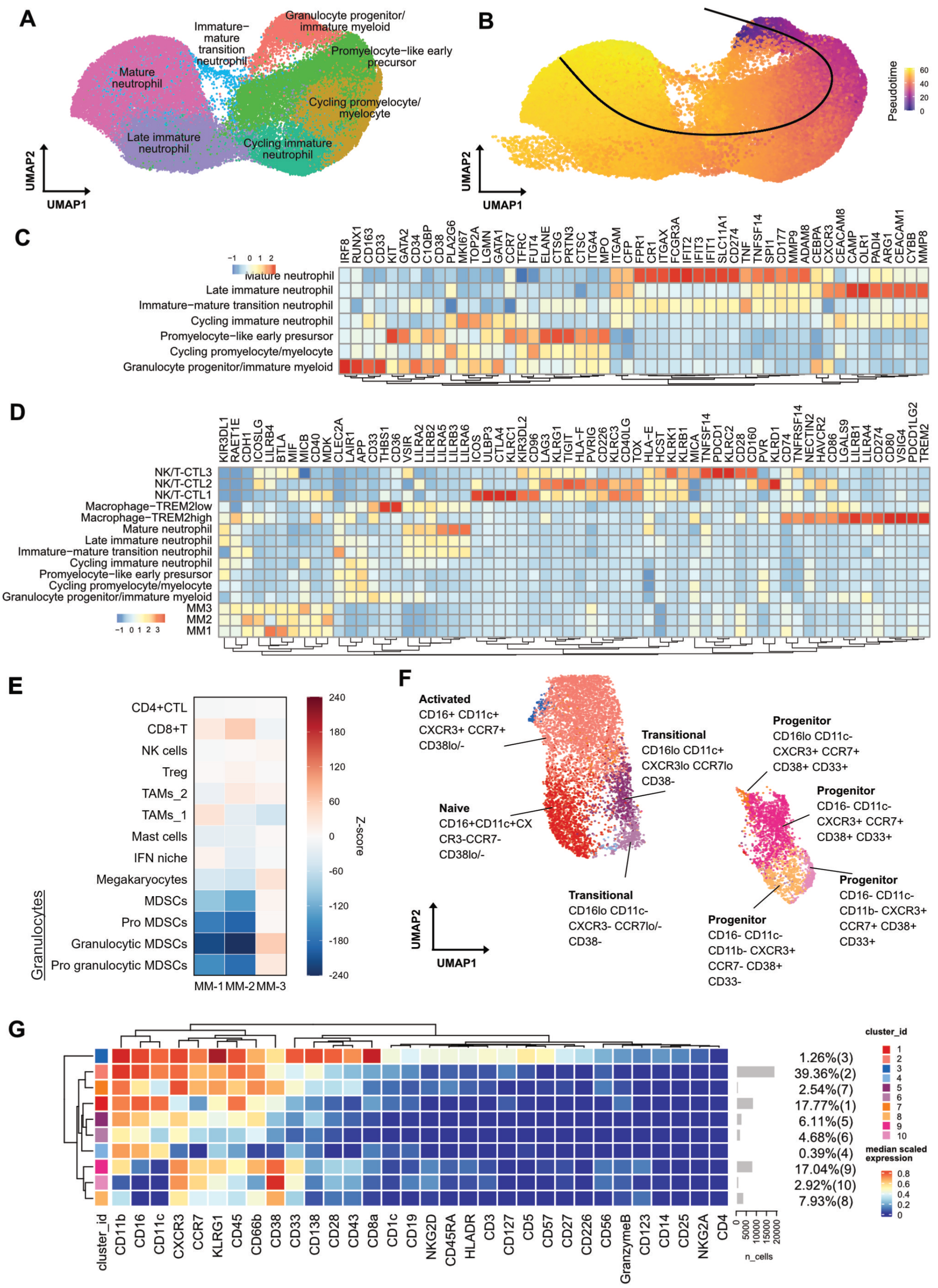

**SI Figure 7. Immune-stromal marker expression of MM1/MM2 regions in Xenium 480 dataset.** **(A)** Heatmap of differential markers between MM1 and MM2 central domain regions (see Figures 4-5 in main text). **(B)** Dotplot for the gene expression profile of functional immune-stromal markers in identified cell populations within central domain regions.

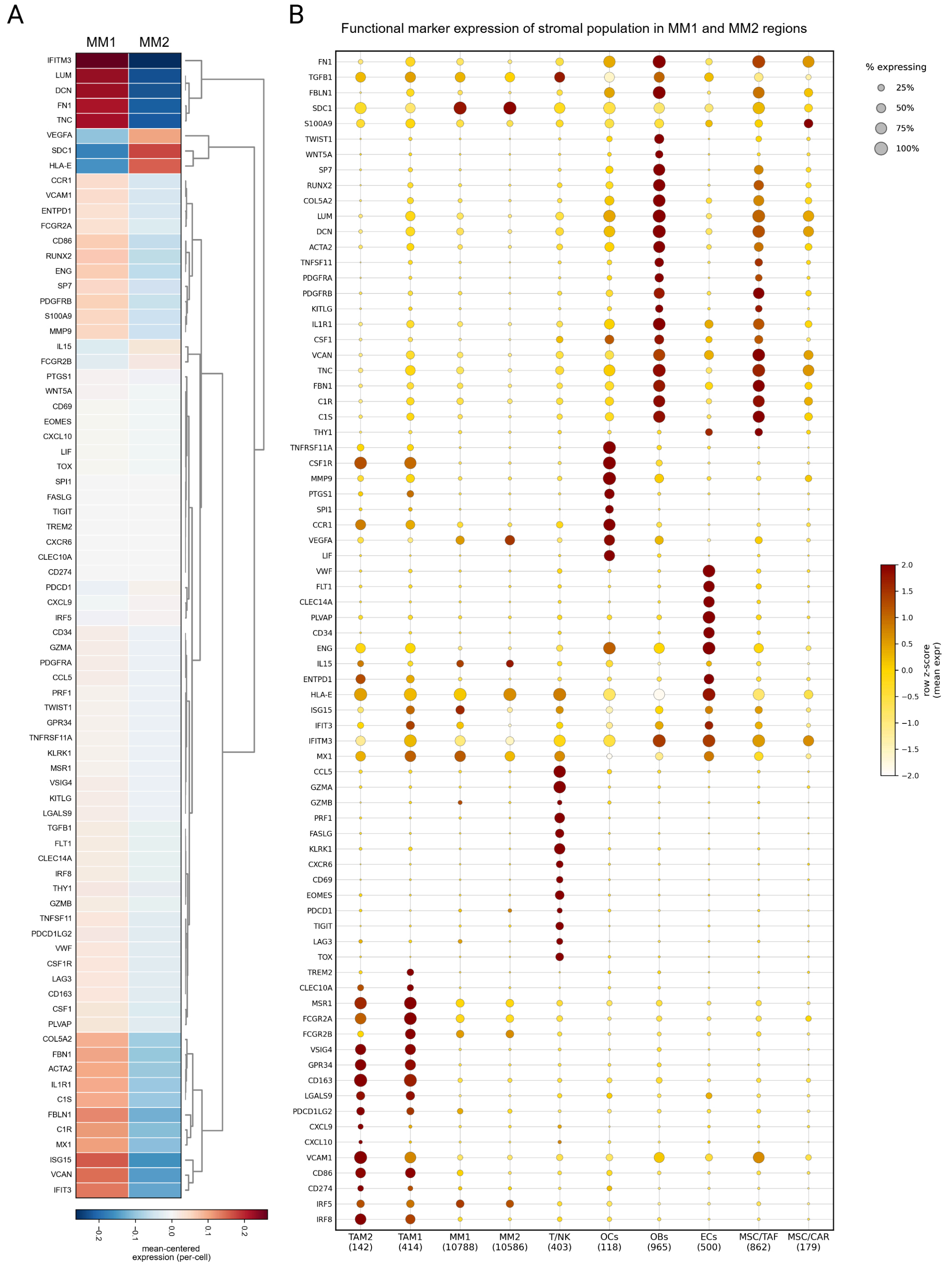

**SI Figure 8. Multi-Omics Factor Analysis (MOFA) integration of spatial WGS, proteomics and pseudobulk RNA data across LCM tiles.** A three modality MOFA model was trained on the 81 laser capture microdissection tiles that passed quality control in all three assays. The model used 4,107 copy number bins for WGS, 5,950 proteins for LC-MS/MS and 480 genes for RNA. Pseudobulk RNA counts were normalised by the total cell number of each tile before modelling. Fifteen factors were fitted. **(A)** Total variance explained by the model in each modality. Bars show the percentage of variance within each modality that is captured by all 15 factors together. **(B)** Variance explained by each individual factor in each modality. Colour shows the percentage of variance in that modality explained by that factor. **(C)** Spatial map of the 81 tiles coloured by MOFA cluster. Clusters were defined by k-means clustering of all 15 MOFA factors with k set to 6. **(D)** Distribution of factor values for Factors 1 to 4 grouped by WGS subclone. Factor 1 separates subclones A and B from subclones C, D and E. Factor 4 separates subclone C from subclones D and E. **(E)** Tiles plotted in MOFA latent factor space. Axes show Factor 2 and Factor 4. Colour shows the MOFA cluster from panel C. Shape shows the WGS subclone. **(F)** The 40 genes with the most negative weights on Factor 4 in the RNA view. **(G)** The 40 proteins with the most negative weights on Factor 4 in the proteomic view.

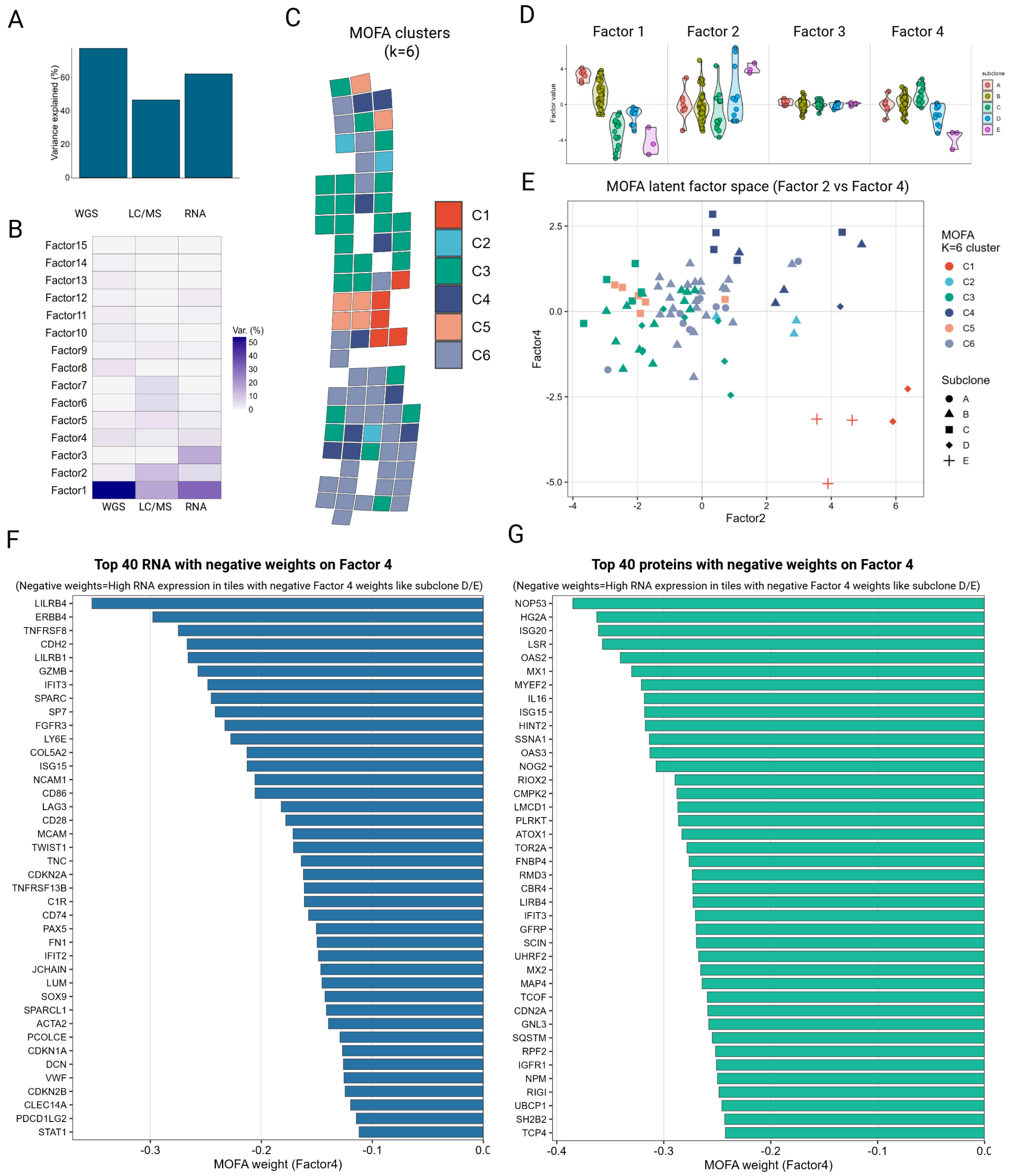
